# Molecular design, optimization and genomic integration of chimeric B cell receptors in murine B cells

**DOI:** 10.1101/516369

**Authors:** Theresa Pesch, Lucia Bonati, William Kelton, Cristina Parola, Roy A Ehling, Lucia Csepregi, Daisuke Kitamura, Sai T Reddy

## Abstract

Immune cell therapies based on the integration of synthetic antigen receptors provide a powerful strategy for the treatment of diverse diseases, most notably retargeting T cells engineered to express chimeric antigen receptors (CAR) for cancer therapy. In addition to T lymphocytes, B lymphocytes may also represent valuable immune cells that can be engineered for therapeutic purposes such as protein replacement therapy or recombinant antibody production. In this article, we report a promising concept for the molecular design, optimization and genomic integration of a novel class of synthetic antigen receptors, chimeric B cell receptors (CBCR). We initially optimized CBCR expression and detection by modifying the extracellular surface tag, the transmembrane regions and intracellular signaling domains. For this purpose, we stably integrated a series of CBCR variants into immortalized B cell hybridomas using CRISPR-Cas9. Subsequently, we developed a reliable and consistent pipeline to precisely introduce cassettes of several kilobases size into the genome of primary murine B cells, again via CRISPR-Cas9 induced HDR. Finally, we were able to show the robust surface expression and antigen recognition of a synthetic CBCR in primary B cells. We anticipate that CBCRs and our approach for engineering primary B cells will be a valuable tool for the advancement of future B cell-based immune therapies.

## Introduction

The successful clinical results of genetically modified T cells for cancer immunotherapy have shown the great potential for engineering immune cells for cellular medicine^1,2,3,4^. Engineered CD8^+^T cells have shown the most progress as they can execute cytotoxic functions by inducing target cells to undergo programmed cell death^5^, thus providing a means to directly attack cancer cells. The strategy to take advantage of the natural features of immune cells, while re-directing their specificity by receptor engineering has culminated in the concept of chimeric antigen receptor (CAR) T cells^6,7,8^. A CAR is a recombinant antigen receptor composed of an extracellular antigen-binding domain, typically an antibody fragment (e.g., a single-chain variable fragment [scFv]), linked by a spacer peptide to a transmembrane domain, which is further fused to an intracellular T cell activation domain, such as CD3ζ^9,10,11^. A broad range of extracellular binding domains and intracellular costimulatory domains (e.g., CD28 and 4-1BB) have been incorporated into CARs to further enhance their targeting and signaling properties^12,13,14,15,16^. CAR T cell therapies rely on the isolation, the *ex vivo* expansion and engineering of T lymphocytes by the introduction of CARs followed by the re-infusion into the patient. While the engineering and development of T cells as cellular therapeutics is advancing rapidly, B lymphocytes represent another class of immune cells that hold promise of being powerful vehicles for adoptive cell therapy due to their involvement in essential processes of immunological recognition and protection. Considering the similarity in the principle of clonal selection and expansion upon antigen exposure, it might be possible to take advantage of natural features of B cells for therapeutic purposes. For example, B cells have very interesting innate properties, such as their ability to differentiate, following antigen-specific activation, into long-lived antibody secreting plasma cells, which home to and reside in specific bone marrow niches, reportedly for decades^17,18^. Their longevity, paired with the capability to secrete large quantities of protein, make primary B cells unique and promising targets as cellular hosts for therapeutic protein production^19^.

Primary T cells can be genetically modified (via lentiviral or retroviral integration) and expanded *in vitro* relatively easily; in contrast, progress on the engineering of B cells has been severely compromised by technical challenges in their *in vitro* culture, expansion and genetic modification. This may be the reason why B cells have received relatively little attention as cellular engineering hosts in immunotherapy. While high rates of transduction in B cells can be obtained using recombinant adenovirus or Epstein-Barr virus vectors, this only results in the temporary expression of transgenes in episomal vectors^20,21^. In contrast, retroviral and lentiviral vectors allow long-term transgene expression by random integration into the host genome. However, these vectors tend to be inefficient at transducing primary B cells^22,23^. Despite the hurldes, there have been a few examples of successful reprogramming of primary B cells: genetically modified B cell have been used for presentation of recombinant antigen for inhibition of immunity in a mouse model of multiple sclerosis^24^ or induction of tolerance towards therapeutic proteins^25^. The revolutionary advances in targeted genome editing have paved the way for alternative strategies to genetically modify immune cells^26,27,28^. So far, the CRISPR-Cas9 system has been mainly applied to integrate transgenes into lymphoma-derived or hybridoma cell lines by homology-directed repair (HDR)^29,30,31^. Precise genome editing in primary murine B cells derived from murine transgenic models endogenously expressing Cas9 protein showed efficient gene disruption through on non-homolgous end-joining (NHEJ) repair^32^. Furthermore, a few recent studies used CRISPR-Cas9 for site-specific gene disruption or transgene integration by HDR in human primary B cells^19,33,34^. Hung *et al.* demonstrated that delivery of Cas9 ribonucleoprotein (RNP) complexes in combination with HDR DNA templates enabled the engineering of plasma cells to secrete therapeutic proteins. This highlights the attractive prospect of establishing a controllable system in which exposure to antigen can induce engineered B cells that produce therapeutic proteins.

Establishing a preclinical genome editing platform based on primary murine B cells enables the investigation of these cells as novel vehicle for adoptive immune cell therapies. In the present study, we have molecularly designed and optimized a novel class of synthetic antigen receptors, chimeric B cell receptors (CBCR), which were stably integrated by CRISPR-Cas9 into immortalized and primary murine B cells. First, we assessed the stable expression of a broad range of constructs encoding a model antigen-specific CBCR linked to a green fluorescent protein (GFP) reporter in a B cell hybridoma line. We genomically modified this cell line by targeting a safe harbor locus (Rosa26) with CRISPR-Cas9 RNP complexes and CBCR HDR templates in the form of linear dsDNA. We then optimized CBCR expression and detection by a series of modifications to the extracellular surface tag, transmembrane domain and intracellular signaling domains. Based on the results obtained from construct screening in hybridoma cells, selected constructs displaying high levels of surface expression were further evaluated in murine primary B cells. Collectively, we could achieve the precise integration of CBCRs into the Rosa26 locus of primary murine B cells, its surface expression and selective enrichment of engineered cells. In the future, CBCR engineered B cells can be evaluated in preclinical *in vivo* models in order to assess their potential in versatile immune cell therapy applications.

## Results

### Design of chimeric B cell receptors (CBCRs)

In this study, we aimed to create a chimeric B cell receptor that is able to recognize antigen independently of the endogenously expressed B cell receptor. We initially used immortalized B cell hybridomas to screen a broad range of CBCR constructs encoding an antigen-binding domain, a spacer region that includes a detection tag, a transmembrane and cytoplasmic signaling domains (**Fig. 1a**). For each of these constructs we generated a stable cell line by CRISPR-Cas9 mediated integration of the transgene cassette into the safe harbor locus Rosa26, which has been validated to stably express robust levels of the transgenes, while minimizing proximity to proto-oncogenes and adverse effects on the host cell. Here, we used a parental hybridoma cell line which constitutively expresses Cas9 from the Rosa26 safe harbor locus approximately 6kb downstream of the CBCR integration site, as it permits the successful editing following transfection of only the pre-formed guide RNA (gRNA, crRNA:tracrRNA complex) and single-stranded oligonucleotide (ssODN) donors^30^. Additionally, this original cell line, that will be referred to as HC9-, contains a frameshift mutation in its endogenous antibody variable heavy chain region, which abrogates antibody expression and makes it a suitable host for the detection of surface-expressed CBCR.

**Figure 1.**
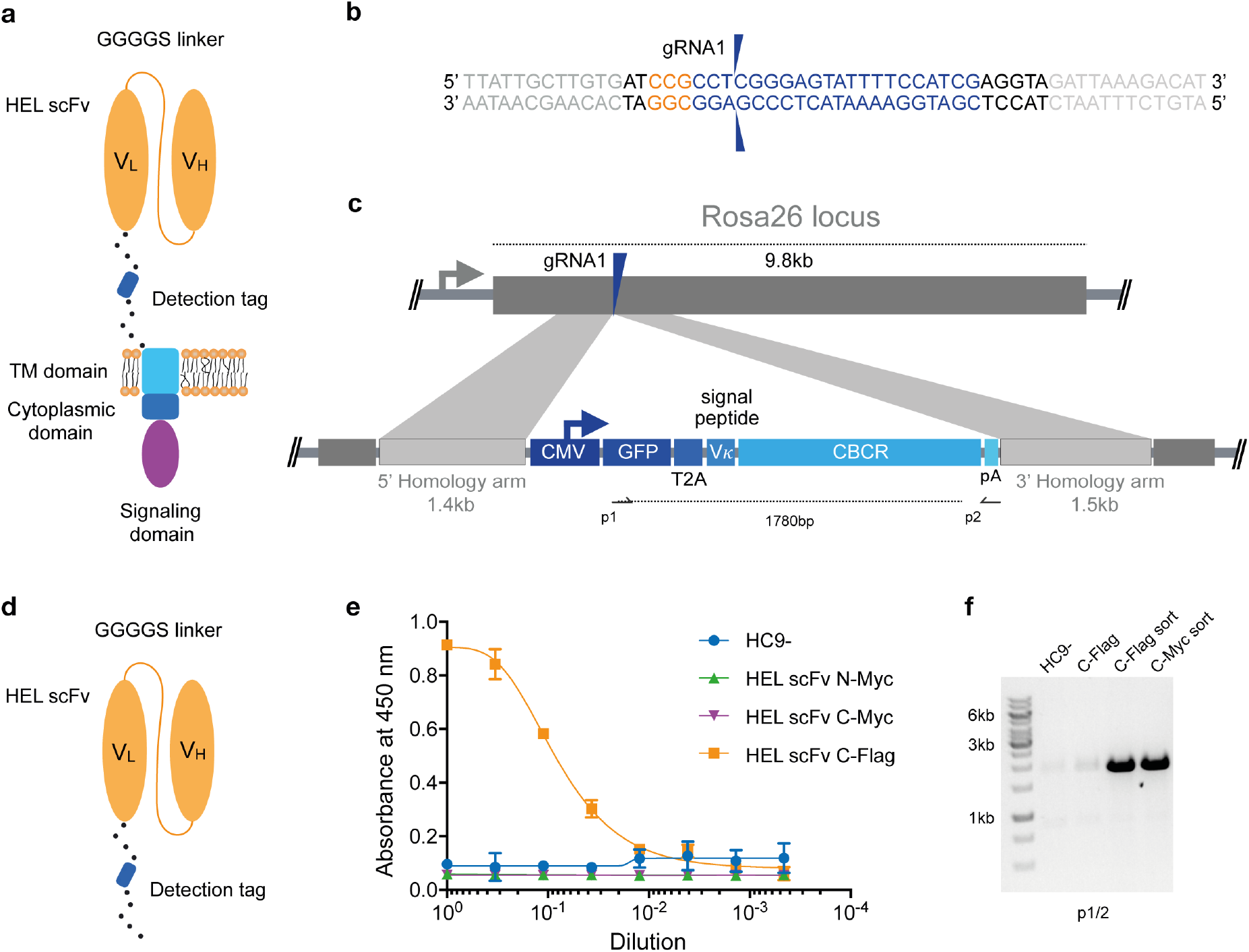
Genomic integration of chimeric B cell receptors (CBCR) by CRISPR-Cas9-mediated HDR. a) Schematic shows the design of the CBCR, which consists of an scFv-based antigen recognition domain (specific for a hen-egg lysozyme [HEL]) (orange), an extracellular spacer with a detection tag, a transmembrane (TM) domain (light blue) and endodomains (blue, purple). scFV: single chain variable fragment, VL: light chain variable domain. VH: heavy chain variable domain. b) Shown is the gRNA sequence (blue) for Cas9-targeting of the safe-harbor locus Rosa26 locus; also shown is the corresponding PAM sequence (orange) as well as the beginning of the homology arms in the HDR template (light grey). The two blue arrows indicate the predicted Cas9 double-stranded break site. c) CRISPR-Cas9-mediated HDR for genomic integration of CBCR construct into the Rosa26 locus. The PCR-linearized donor template contains a GFP reporter gene followed by a T2A coding sequence, the *V_κ_* signal peptide, the CBCR cassette and a polyA sequence under control of a CMV promoter and flanked by sequences homologous to the Rosa26 locus next to the gRNA target site of 1.4kb and 1.5kb, respectively (light purple). T2A: the self-cleaving thoseaasigna virus 2A sequence, pA: SV40 polyA sequence d) Schematic shows the CBCR antigen binding domain including a linker with a detection tag, either Myc or Flag epitope. e) Cells were enriched for GFP (488 nm) expression via FACS. Graph shows ELISA results of scFv secretion levels (capture HEL antigen, detection anti-Myc or Flag) on enriched hybridoma culture supernatant for scFv variants (excluding the TM and intracellular domains shown in d) with Myc or Flag detection tag in N- or C-terminal position. Supernatant of HC9-cells was used as negative control. For each sample, three technical replicates were analyzed and a four-parameter logistical curve was fitted to the data by nonlinear regression. Data are presented as the mean and error bars indicate standard deviation. f) RT-PCR on mRNA of scFv variants with C-terminal detection tags was performed with primers shown in c displaying expression of the transgenic scFv cassette after transfection only and transfection followed by sorting on GFP expression.

Within the 5’ portion of the Rosa26 locus, we identified several potential gRNA sites compatible with *S. Pyogenes* Cas9 and its protospacer adjacent motif (PAM, 5’-NGG). The activity of Cas9 at each gRNA site in B cells was evaluated by measuring non-homologous end-joining (NHEJ) via Surveyor nuclease assay (Supplementary **Fig. S1**)^35^. The gRNA with the highest activity (gRNA1) was selected for all subsequent genome editing experiments (**Fig. 1b**). To precisely integrate the CBCR by Cas9-mediated HDR, donor templates were designed including the respective CBCR transgene flanked by homology arms of 1.4kb/1.5kb length complementary the Rosa26 sequences next to the gRNA1 target site (**Fig. 1c**). The PAM was not incorporated in the repair template, so that the repair template and the repaired sequence would not to be cleaved by Cas9. The full transgene consisted of a V_κ_ leader sequence, a GFP reporter gene followed by a self-cleaving T2A sequence, and the CBCR all under the control of the cytomegalovirus (CMV) promoter^36^. The HDR donor was generated by PCR to obtain a linearized format and electroporated alongside the gRNA1 into the HC9-hybridoma cells. At ~72 h post-transfection, GFP^+^ cells were isolated by fluorescence-activated cell sorting (FACS) and expanded in culture.

For the extracellular antigen-binding domain, we used scFvs, as these have been successfully used in previously engineered receptors such as CARs and synNotch^10,37^. Additionally, scFvs offer great stability and high-affinity ligand binding^38^. As a target, we selected the model protein hen egg lysozyme (HEL) due to its small size, easy availability and the presence of valuable research tools such as a HEL-specific B cell transgenic mouse model and well-described HEL-specific antibodies and scFvs^39,40,41^. Initially, we designed an scFv from the high-affinity antibody HyHEL10 in the V_L_-V_H_ orientation. After comparison of this scFv to the affinity-improved M3 mutant scFv derived from the anti-HEL antibody D1.3, we proceeded with the M3 scFv in the V_H_-V_L_ orientation due to increased detection signal by enzyme-linked immunoabsorbent assays (ELISA) (**Fig. S2**)^42^. CBCR expression can be detected by antigen labeling, however, this is often accompanied by B cell activation, therefore an orthogonal detection method would be valuable. While the GFP offers a selection marker for integration, it does not directly indicate surface expression of CBCR, thus the tactical introduction of a detection tag provides another identification marker. As has been previously shown with CARS in T cells, careful design of tag sequences and their location is required^43,44,45^. Initially the M3 scFv was equipped with an N-terminal Myc epitope (EQKLISEEDL) or fused to a C-terminal spacer sequence (26 amino acids [aa]) incorporating a Myc or Flag epitope (DYKDDDDK) (**Fig. 1d**). We used a secretion variant of the CBCR, which lacks the transmembrane and intracellular signaling domains to evaluate integration and secretion levels of HEL-binding scFv. We used enzyme-linked immunosorbent assays (ELISAs) on normalized culture supernatants.

Drastic improvement in scFv secretion was observed for cells in which the Flag sequence was introduced into the C-terminal spacer as compared to both Myc epitope containing variants (**Fig. 1e**), whereas equal RNA expression levels were confirmed by RT-PCR (**Fig. 1f**) indicating impairment of either proper scFv protein folding or secretion by the Myc epitope tag. RNA expression of C-terminally incorporated Flag tag was significantly increased after enrichment for GFP^+^ cells via FACS compared to unsorted cells (**Fig. 1f**).

### Optimization of CBCR for robust surface expression on hybridoma B cells

Next, we investigated whether a CBCR with the previously characterized scFv domain was presented at the cell surface, while maintaining antigen (HEL) binding. For this purpose, we linked the scFv clone M3 and the spacer incorporating the Flag epitope sequence to the transmembrane domain (TM) and the short cytoplasmic domain of the endogenous murine BCR (IgG2c) referred to as CBCR-BCR-TM. Alternatively, the spacer was fused to a CD28 transmembrane domain (CBCR-CD28-TM), which has been successfully incorporated in the design of several CARs^46^ (**Fig. 2a**). When expressed in HC9-hybridoma cells, both CBCR-BCR-TM and CBCR-CD28-TM were detected on the cell surface based on HEL antigen binding for cells enriched for GFP by FACS (**Fig. 2b**, upper row). CBCR-CD28-TM was expressed at the cell surface to a substantially greater extent than CBCR-BCR-TM. Additionally, CBCR cells were stained with anti-Flag antibody. The CBCR surface expression was demonstrated by HEL antigen recognition in GFP^+^ cells, but this did not correlate with the CBCR detection via the Flag peptide tag, suggesting impaired accessibility of the Flag epitope, once the spacer is fused to a transmembrane domain (**Fig. 2b**, lower row). However, we were able to identify clones demonstrating both a clear Flag tag expression and HEL antigen binding after performing single-cell sorting and expansion (**Fig. 2c**, right). While exhibiting a similar level of stable GFP expression (**Fig. 2c**, left), CBCR-CD28-TM surface expression was increased compared to CBCR-BCR-TM expression on the surface, consistent with the data obtained from bulk-sorted cells detected by HEL binding only (**Fig. 2b**). To evaluate stable and targeted integration of the CBCR cassette on a genotypic level, PCR assays on genomic DNA were designed (**Fig. 2d**). The introduced cassette was detected by end-point PCR in the single-cell line expressing CBCR-BCR-TM, but not in the parental HC9-cell line (primers p3/4). PCR analysis showed the presence of at least one residual wildtype allele in the cell line (**Fig. 2e**). Genomic PCR using primers p5 and p6 confirmed precise integration of the CBCR gene into the Rosa26 locus (**Fig. 2f**).

**Figure 2.**
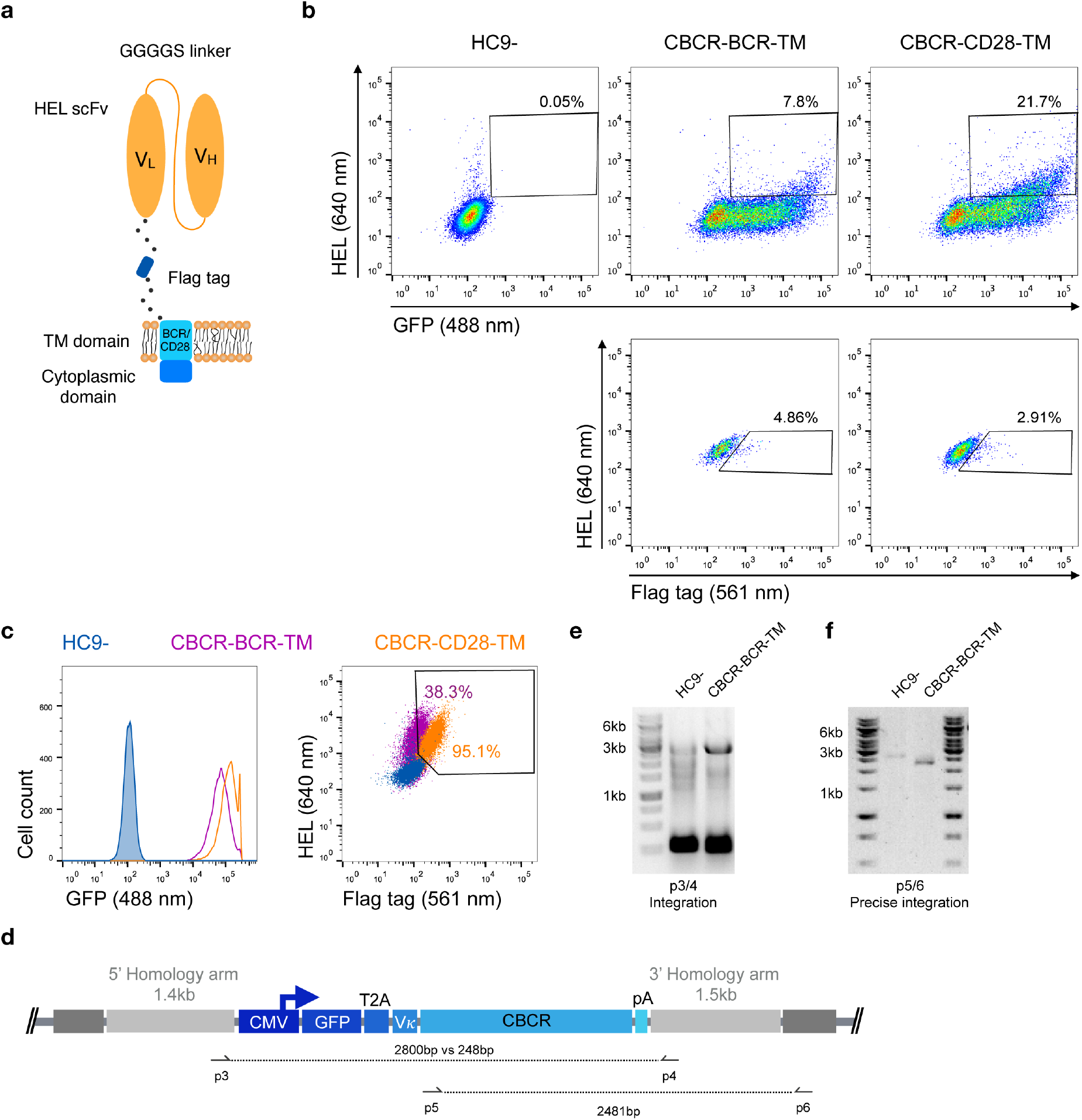
Stable surface expression of CBCR on hybridoma cells. a) Schematic of surface expressed CBCR including a Flag detection tag and varying transmembrane (TM) domains. The TM domain is derived from either the endogenous BCR composed of an immunoglobulin (purple in c) or the T cell costimulatory CD28 molecule (orange in c). b) Representative flow cytometry dot plots of the cells after enrichment based on GFP reveal CBCR surface expression (via HEL antigen binding) for both variants (upper row), but no correlation with surface detection via the Flag protein tag (561 nm, lower row). The parental hybridoma cells were used as negative control. Cells that were positive for HEL binding and Flag tag expression were enriched by FACS. c) Flow cytometry analysis of resulting single-cell clones selected for GFP expression and binding to HEL shows comparable levels of persistent GFP expression for both construct variants (left), but differs in terms of CBCR surface expression (right). d) Schematic of primer sets for genomic DNA analysis in order to detect transgene integration (p3/4) and to confirm GFP-2A-CBCR cassette integration at the correct Rosa26 locus (p5/6). e) Agarose gel shows genomic PCR products that confirm the presence of the transgene in the cell line expressing the CBCR containing the BCR transmembrane (2800bp). Additionally, the presence of at least one wt allele is demonstrated by the PCR product with a length of 248bp. f) Genomic PCR analysis verifies the integration of the GFP-T2A-CBCR-BCR-TM cassette in the correct locus (2481bp) in the same cell line as in e. PCR products in e and f were verified by Sanger sequencing.

### Strep tag II incorporation improves CBCR surface expression and selection

Previous studies have reported that the length of the non-signaling extracellular spacer can have an impact on surface expression or receptor activity^37,45^. We constructed variants with shorter and extended spacers in combination with both TM domains in order to analyze the influence on surface expression of the CBCR and the accessibility of the Flag detection tag within the spacer. HEL antigen binding within the GFP^+^ population was examined for cells expressing CBCR including spacer regions of different lengths before and after sorting of GFP^+^ cells (**Fig. 3a and b**). Antigen binding did not vary significantly with extracellular linker length, indicating that the composition of the spacer does not affect the surface expression in these cases. Detection of the Flag tag in GFP^+^ bulk-sorted cells still was not improved after modifying the length of the spacer (**Fig. 3c**). To address the tag detection, we introduced one or more Strep tag II sequences replacing the Flag epitope within the Gly/Ser spacer (**Fig. 3d**). All Strep tag CBCR were stained with anti-Strep tag II antibody after enrichment of GFP^+^ cells via FACS. Staining intensity was significantly increased for both CBCR-TM variants with one Strep tag II compared to the signal provided by the single Flag epitope (**Fig. 3e**). Surface expression based on antigen binding and tag detection again revealed increased level for CBCR-CD28-TM compared to the CBCR-BCR-TM variant. Further, staining intensity was dramatically enhanced for CBCR-CD28-TM cells that contained three Strep tag II sequences (**Fig. 3e**, right). This data indicate that inclusion of Strep tag II improves the CBCR surface expression and its correlation with staining based on the detection tag.

**Figure 3.**
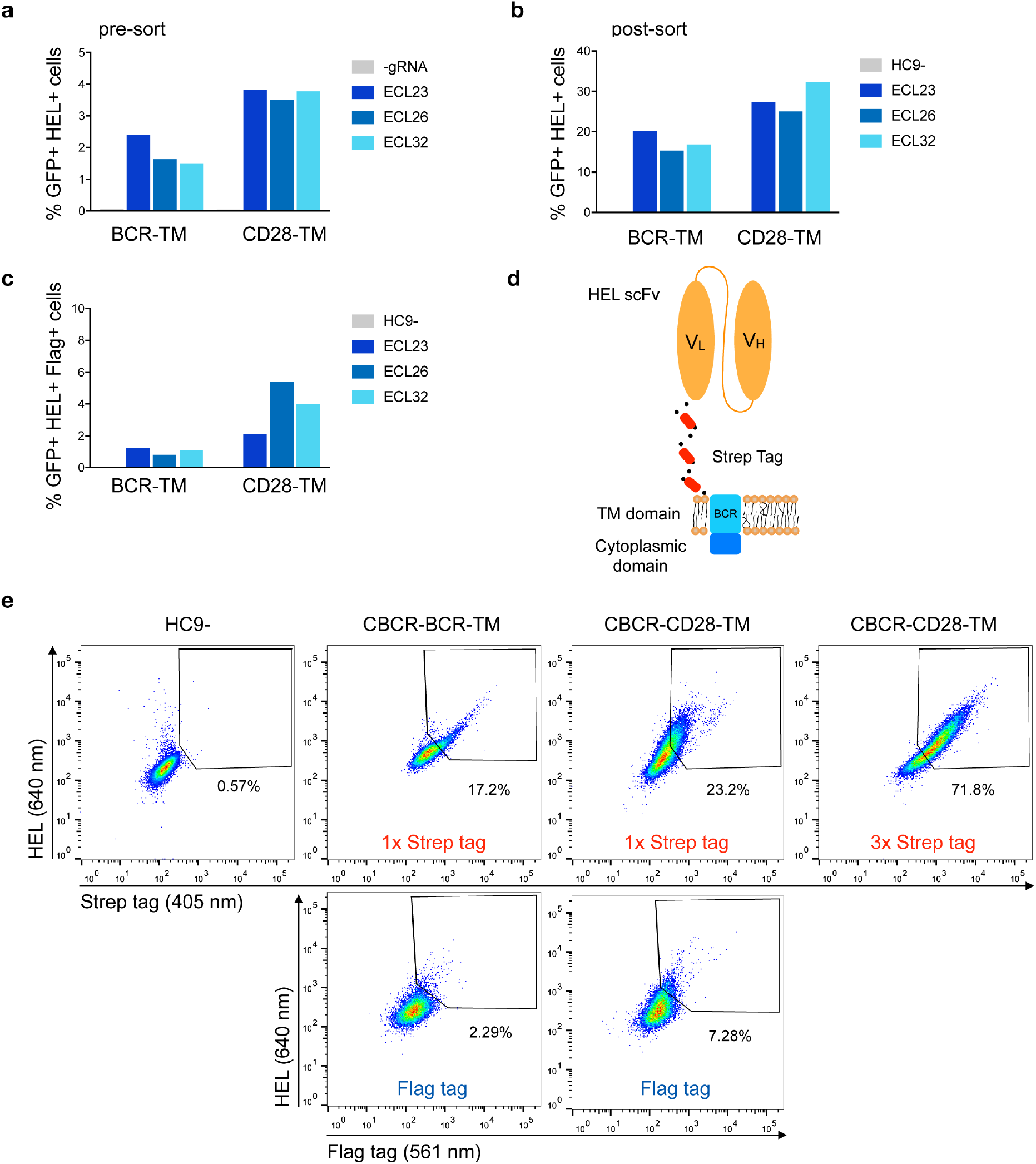
Optimization of CBCR receptor design to improve surface expression and the detection of surface presentation via tag. a) The surface expression in hybridomas of representative CBCR constructs was evaluated by flow cytometry before (a) and after (b) enrichment based on GFP expression (488 nm) on day 3 after transfection. Variants shown differ in the transmembrane (TM) domain and the lengths of the extracellular linkers (ECL). c) Bar graph shows percentage of cells detectable via Flag tag within the population (b) of cells showing GFP^+^ expression and HEL binding after sorting. d) Schematic of CBCR design shows replacement of Flag epitope with a triple Strep epitope II tag (red). e) CBCR surface expression of GFP sorted cells (488 nm) was quantified by flow cytometry based on Strep II tag detection (405 nm) using variants that incorporate a single or triplet Strep epitope (upper row). Antigen recognition was additionally confirmed by HEL binding (640 nm). Representative flow cytometry plots show percentages of HEL binding and Flag^+^ (561 nm) cells for the CBCR variants including the single Flag tag after GFP sorting as a control (lower row).

### Incorporation of a CD79b signaling domain improves CBCR surface expression

In order to generate a functional receptor, an intracellular signaling domain is required for signal transduction. As the endodomain of surface-bound immunoglobulins (Ig) itself is very short and incapable of intracellular signal transmission, the endogenous BCR only functions as a complex composed of the Ig molecule associated with a heterodimer called Ig-α/Ig-β or CD79a/b. The CD79 subunits contain two immunoreceptor tyrosine-based activation motif (ITAMs) in their cytoplasmic domains, which recruit Syk tyrosine kinase and mediate B cell activation upon antigen binding and subsequent phosphorylation^47,48^. Previous work suggested that CD79a and CD79b are each independently sufficient to trigger protein tyrosine kinase activation and induction of downstream signaling cascades, as long as the ITAM regions remain intact^49^. Here, we fused the complete intracellular domain of either the CD79a or CD79b polypeptide to the short cytoplasmic tail of the Ig molecule at the C-terminal end (**Fig. 4a**). HC9-cells were transfected with both constructs as previously described and stable CBCR expression based on HEL binding and Strep tag II detection was analyzed using flow cytometry after enrichment for GFP^+^ cells. CBCR expression was substantially increased for cells expressing CBCR-CD79b compared to CBCR-CD79a, which was however higher compared to previous constructs without intracellular signaling units (**Fig. 3e**, upper row). This suggests that improved expression or cell-surface transport for the CBCR variant occurs when incorporating the CD79b signaling domain (**Fig. 4b**). This higher CBCR expression was consistent across several selected single-cell clones (**Fig. 4c**, right), while GFP expression was comparable, but intensity was slightly increased for CBCR-CD79b expressing cells (**Fig. 4c**, left). Genomic PCR analysis using primers p7 and p8 verified precise integration of the GFP-CBCR-CD79b cassette into the Rosa26 locus in one of the selected single-cell clones (**Fig. 4d and e**). Conclusively, incorporation of a signaling domain did not interfere with CBCR surface expression.

**Figure 4.**
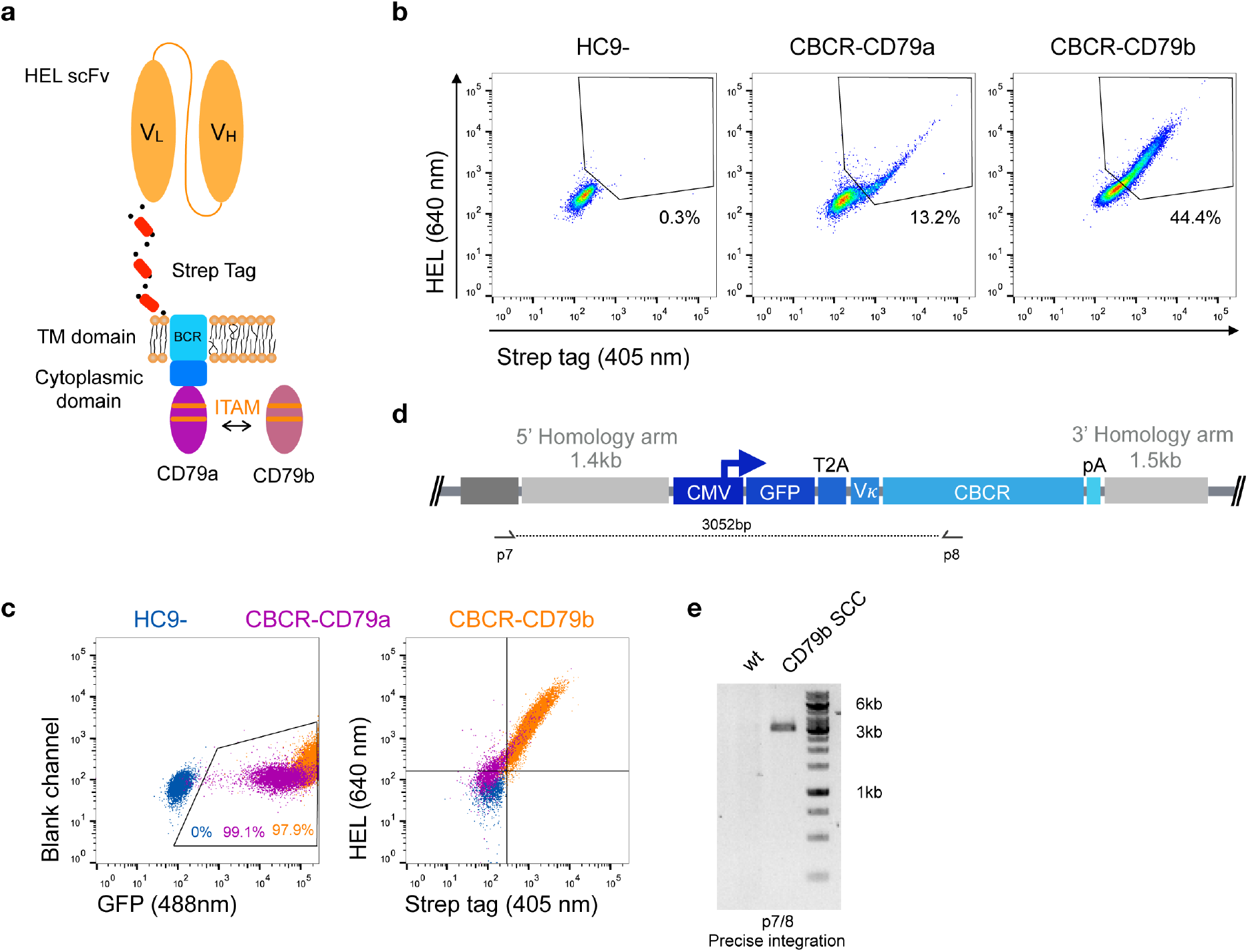
Improved surface expression of CBCR with CD79b signaling domain. a) Schematic of the CBCR complex containing the intracellular domain of either a CD79a or CD79b transmembrane protein C-terminally fused to the cytoplasmic BCR domain. b) Representative flow cytometry dot plots show hybridomas with stable CBCR surface expression based on HEL binding (640 nm) and Strep II tag detection (405 nm) for GFP enriched cells having a CBCR with either the CD79a or CD79b intracellular domain precisely integrated. Data are representative for three independent experiments. c) Flow cytometry dot plots show GFP expression (left) and HEL binding and Strep II tag detection of exemplary single-cell clones following sorting (HEL^+^ Strep^+^) on samples in b. Tendency of decreased surface expression for the CBCR with the CD79b intracellular domain was validated in multiple single-cell clones. d) Schematic of primer set for genomic DNA analysis in order to confirm integration of GFP-2A-CBCR cassette at the correct locus (p7/8). e) Genomic PCR analysis verifies the integration of the GFP-T2A-CBCR-CD79b cassette into the correct locus of a single-cell clone (SCC, 3052bp). The band was extracted and Sanger sequencing confirmed the precise integration in the Rosa26 locus.

### Evaluating HDR protocols for primary murine B cells

High genome editing efficiencies are not essential when engineering hybridoma cells, which have the capacity to expand at high rates and can be cultured without time constraints. However, with primary B cells, high HDR efficiencies are very important. We first isolated murine splenic B cells and cultured them using an *in vitro* expansion system based on 40LB feeder cells (Balb/c 3/T3 fibroblasts that stably express BAFF and CD40 ligand)^50^ in the presence of IL-4 up to four days for pre-activation (**Fig. 5a, c**). Next, we electroporated Cas9-guide RNP complexes together with 5 μg of PCR-linearized HDR donor template into primary B cells. Three days after transfection, cells were analyzed and enriched for GFP expression by FACS. Sorted cells were cultured for an additional six days under activating conditions by replacing IL-4 with IL-21. CBCR surface expression was determined via flow cytometry and genomic DNA analysis was performed at Day 10 to confirm precise integration into the Rosa26 locus (**Fig. 5a**). Overall increase in the number of live B cells on 40LB feeder cells in presence of IL-4 was only negligibly affected by transfection of Cas9-RNP and dsDNA as compared to non-transfected cells co-cultured (**Fig. 5b**). In contrast, primary B cells cultured in the presence of soluble BAFF and IL-4 showed only a minor level of expansion. (**Fig. 5b**). Subsequently, the influence of pre-activation on transfection efficiencies was determined. For this purpose, we transfected 10^6^ primary B cells with 2 μg plasmid DNA encoding for a CMV promoter-driven GFP reporter gene directly after isolation from a mouse spleen or following pre-activation on 40LB feeder cells for one, two, three, or four days and analyzed GFP expression by flow cytometry 24 h after transfection. We observed enhanced transfection efficiencies and viability after pre-activation compared to transfection of freshly isolated B cells, consistent with previous findings in primary T cells^51^. The highest transfection efficiency was observed after one day of pre-activation, followed by four days of pre-activation, suggesting a correlation with the time points of high proliferation rates. Finally, a protocol was designed to integrate the best condition for transfection efficiency, one day of pre-activation, with delivery of components previously optimized in hybridoma cells; however, only very small numbers of precisely edited primary cells, identified by GFP expression, could be obtained with this workflow (0.1-0.3%).

**Figure 5.**
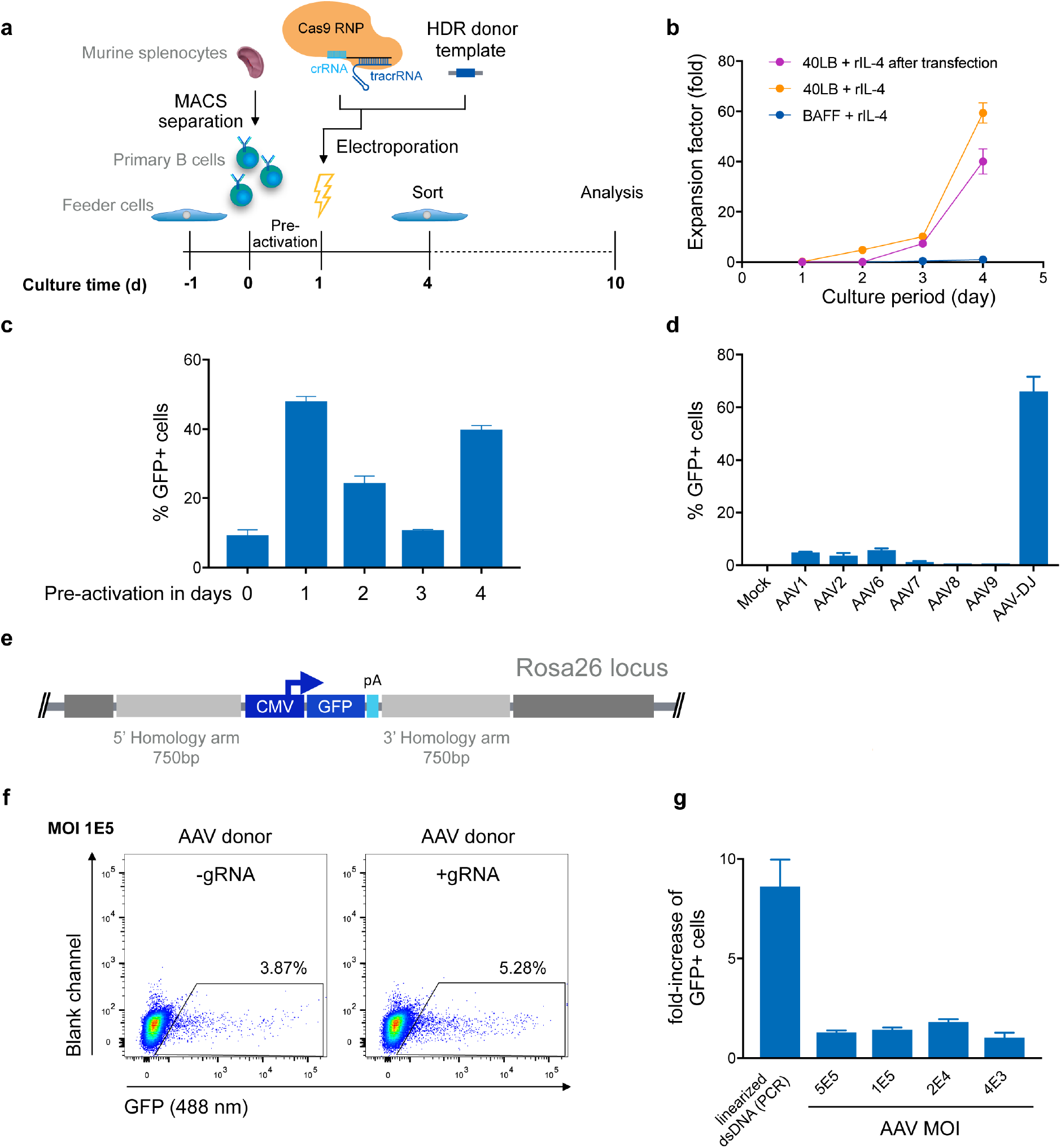
Targeted genome editing in primary murine B cells using CRISPR-Cas9. a) Overview shows timeline of Cas9-gRNA RNP delivery to primary B cells isolated from murine spleen. B cells were initially co-cultured with feeder cells before and after transfection of RNPs and HDR donor template DNA on day 1. Three days after transfection cells were enriched for GFP expression via FACS and analyzed at Day 10 after expansion. b) Cumulative fold increase in the number of live B cells cultured in the following conditions: (i) in presence of soluble BAFF and IL-4; (ii) on 40LB cells in the presence of IL-4, (iii) or on 40LB in the presence of IL-4 and after electroporation with Cas9-RNP and PCR-linearized double-stranded (ds) repair template DNA after pre-activation on 40LB for one day. c) 10^6^ primary B cells were transfected with 2 μg plasmid DNA (pMax-GFP) directly after isolation from mouse spleen or following pre-activation on 40LB feeder cells for one, two, three or four days. Data show the percentage of GFP expressing cells determined by flow cytometry 24 h after transfection. Data are presented as the mean and error bars indicate standard deviation (n=2). d) Splenic B cells were preactivated for one day and were either mock treated or transduced with GFP-expressing ssAAV using a comprehensive panel of AAV serotypes (1, 2, 6, 7, 8, 9, or DJ) at a MOI of 10^5^. The bar plot shows the percentages of GFP^+^ cells after 72h (n=3, 3 independent experiments). e) Schematic of the HDR donor cassette encoding for the GFP reporter gene with 750bp flanking homology arms after integration into the Rosa26 locus. f) After one day of pre-activation on 40LB cells, primary B cells were transfected with Cas9-RNP immediately followed by HDR repair template delivery via chimeric AAV serotype DJ encoding the GFP reporter gene. Representative flow cytometry dot plots show GFP expression (488 nm) on day 9 after genome editing for transduction with a MOI of 1 x 10^5^. Cells transfected only with Cas9-protein without gRNA and transduced with GFP expressing HDR donor packaged using scAAV-DJ were used as negative control to determine the level of GFP expression from episomal retention. g) Data are displayed as fold-increase of AAV-DJ transduced GFP^+^ cells receiving the Cas9-gRNA complex to cells transfected with Cas9-protein only representing the HDR based integration. Cells transfected with Cas9-protein only indicate the episomal AAV background expression. Cells transfected with Cas9-RNP and PCR-linearized dsDNA served as control. GFP expression was measured on day 9 after transfection. All data are means ±s.d (n=3).

Several studies suggest that the use of single-stranded (ss) DNA increases the frequency of HDR, most notably through the use of adeno-associated virus (AAV)^52,53,30^. AAV can package a genome of at least 4.9kb, a length sufficient for an HDR donor template compatible with CBCR and homology arms. Previous studies have reported relatively high levels of AAV-mediated HDR in multiple cell types, including T lymphocytes^54,55,56^. To investigate AAV transduction efficiency in primary murine B cells, we screened several AAV serotypes possessing a reporter GFP gene (**Fig. 5d**). We transduced B cells after pre-activation on 40LB for one day and analyzed transient GFP expression by flow cytometry after three days, which represented the overall transduction efficiency. The highest transduction efficiency was achieved with the synthetic AAV-DJ serotype (**Fig. 5d**). Regardless of serotype, we observed minimal loss in cell viability after exposure to the virus particles. Next, we examined the frequency of HDR-mediated integration of a larger size transgene delivered by recombinant AAV-DJ. For this purpose, an HDR donor cassette consisting of a CMV-driven GFP reporter gene was designed with homology arms of 750bp in size to meet AAV payload restrictions (**Fig. 5e**). After pre-activation and electroporation with or without complete Cas9-RNP, B cells were transduced with AAV-DJ CMV-GFP at various multiplicity of infections (MOIs) and cultured for an additional six days in the presence of 40LB cells and IL-21 (**Fig. 5f and g**). We observed only minor loss in viability, even at the highest AAV dose. Approximately 3-4% of cells that were treated with AAV alone (MOI 1 x 10^5^) showed persistent GFP expression implying a relatively high background of episomal expression (**Fig. 5f**). In cells that received both AAV-DJ delivered HDR donor and Cas9-RNP (targeting the Rosa26) we observed only a marginal increase in HDR (1.2-1.5-fold), measured by stable GFP expression. When these conditions are compared with cells receiving Cas9-RNP and an HDR template in the format of PCR-linearized dsDNA versus PCR-derived template DNA only, we observed up to a ten-fold increase in HDR efficiency (**Fig. 5g**). While HDR efficiencies in primary B cells were only marginally enhanced by AAV-DJ delivered HDR donor, HDR-mediated integration efficiencies were dramatically improved in hybridoma cells, suggesting that primary B cells may have inherent limitations in HDR processes (**Fig. S3**). When directly compared to dsDNA, we found that AAV-DJ showed slightly improved HDR-mediated integration, however it also resulted in strong background GFP expression complicating the discrimination of precisely edited cells especially considering the limited life span and restrictions in selection of primary B cells in *in vitro* culture (**Fig. 5g**). The results described here suggest that despite the relatively low HDR-efficiencies, dsDNA HDR template is more suitable for CRISPR-Cas9-mediated genome editing in primary B cells due to its reliable discrimination of successfully modified cells.

### Robust CBCR genomic integration and surface expression of CBCR in primary B cells

We evaluated the surface expression of the previously optimized CBCR variants in order to generate primary B cells capable of antigen recognition independent of their endogenously expressed BCR. For this purpose, we transfected pre-activated B cells with Cas9-RNP targeting the Rosa26 locus and PCR-derived HDR donor (GFP-T2A-CBCR-CD79a/b and GFP-T2A-CBCR-BCRTM/CD28TM) and incubated them on 40LB feeder cells in the presence of IL-4 for recovery. At day three after electroporation, we observed low, but robust HDR-mediated integration levels, measured by persistent GFP expression, compared to a negative control of cells receiving PCR-linearized repair template and Cas9 protein without gRNA (**Fig. 6a**, upper row). GFP^+^ cells were enriched via FACS, expanded in the presence of IL-21 and analyzed by flow cytometry for CBCR surface expression based on HEL antigen binding and Strep tag II detection on day 10 (**Fig. 6b**). We found substantial enrichment for GFP expressing cells, from which a robust fraction is expressing either CBCR variant, thus, indicating that CBCR expression is tolerated in primary B cells. HEL antigen binding by the CBCR does not appear to be inhibited by expression of native BCR. Similar to our observations in hybridoma B cells, CBCR detection based on the Flag tag was rather weak (**Fig. 6b, S4a**). Consistent with hybridomas, an enhanced CBCR-CD79b expression compared to CBCR-CD79a was confirmed in primary B cells. PCR analysis on genomic DNA extracted from primary B cells with or without FACS mediated enrichment verified the targeted integration of the GFP-T2A-CBCR cassette into the Rosa26 locus (**Fig. 6c, S4b**). Taken together, these findings demonstrate that we have developed a reliable and consistent pipeline to precisely introduce cassettes of several kb size into the genome of primary murine B cells using CRISPR-Cas9 induced HDR. Furthermore, we were able to show the robust surface expression of a synthetic, antigen-specific CBCR in primary B cells.

**Figure 6.**
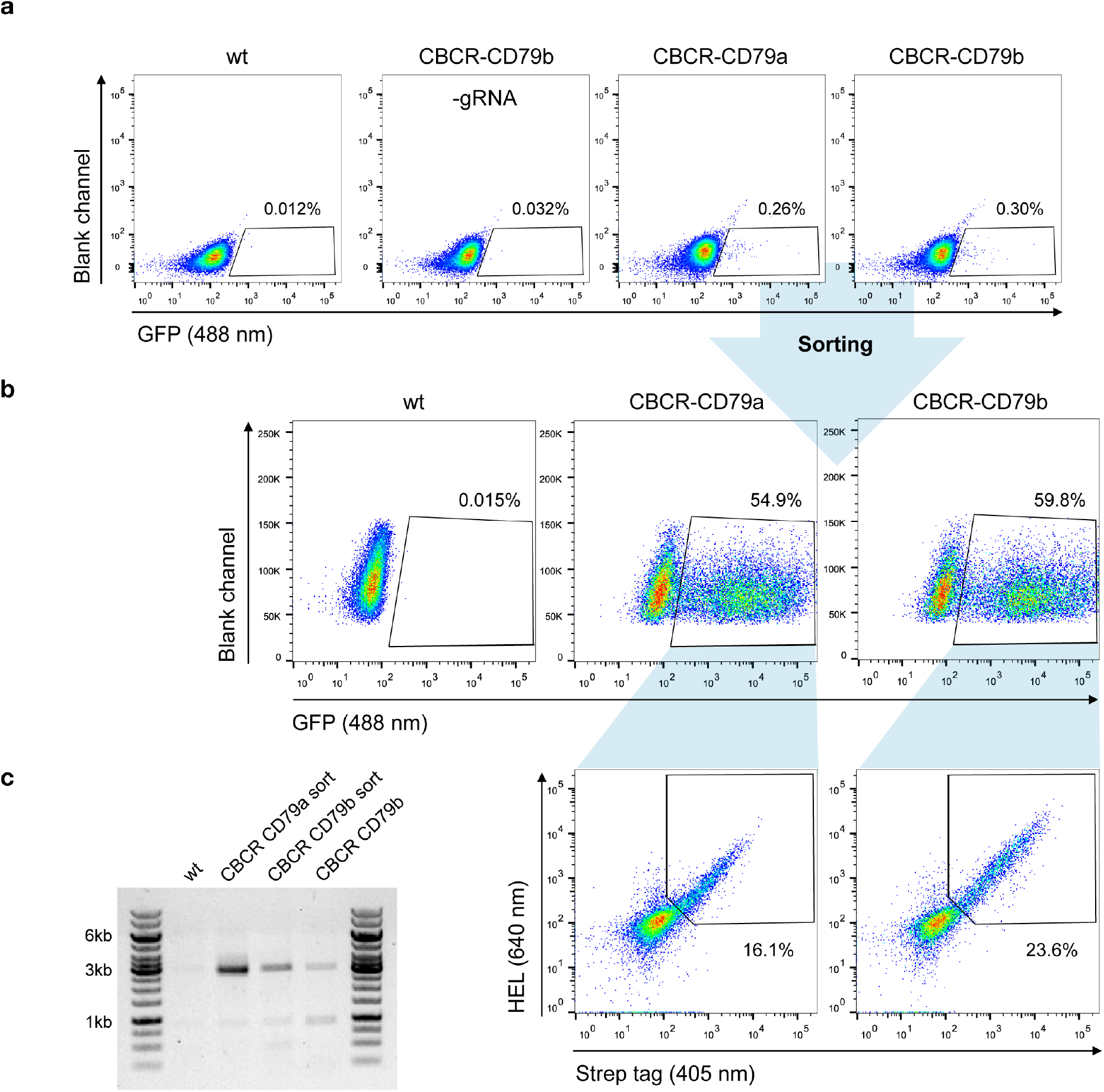
Robust and stable CBCR surface expression in primary B cells. a) Splenic B cells from C57BL/6-Ly5.1 mice were transfected with Cas9-RNP and HDR donor templates encoding synthetic CBCR (previously optimized in hybridoma cells, Figure 3 and 4) following 24 h of pre-activation on a 40LB feeder cell layer and cultured in the presence of IL-4. Integration efficiencies based on GFP expression (488 nm) were determined by flow cytometry on day 3 after transfection and GFP^+^ cells were sorted. Primary B cells electroplated without gRNA and non-transfected B cells serve as negative controls. Flow cytometry plots are representative of three independent experiments. b) Sorted primary B cells were successfully regenerated during co-culture on 40LB feeder cells and in the presence of IL-21. Flow cytometry dot plots show efficient enrichment of GFP^+^ cells (upper row) and CBCR surface expression in primary B cells based on HEL antigen binding and detection of the Strep II tag within the GFP^+^ population (lower row). c) Agarose gel shows genomic PCR products (p7/8) that confirm the targeted integration of the GFP-T2A-CBCR cassette containing the intracellular domain of either CD79 protein (3052bp) before and after sorting.

## Discussion

Immune cell therapies based on the integration of synthetic antigen receptors comprise a successful and rapidly expanding therapeutic option for the treatment of cancer, most notably CAR expressing T cell therapies^1,37,45^. Additional to existing T cell therapies, B lymphocytes hold promise as novel donor cells for adoptive cell therapies due to their natural properties, such as longevity and immense protein secretory capacity^18,19^. Here, we have demonstrated targeted genomic integration in murine B cells of a novel class of synthetic antigen receptors, CBCR. CBCRs offer a potential way to activate and expand engineered B cells in antigen-controllable manner, independent of the endogenously expressed BCR.

We designed CBCR constructs to encode an antigen-binding domain consisting of an scFv, a spacer region that includes a detection tag, a transmembrane domain and cytoplasmic signaling domains. Detection tags incorporated into the extracellular spacer provide a valuable identification marker for receptor surface expression. We observed dramatic differences in secretion and detection of surface expression levels for the analyzed constructs (**Fig. 1**). Interestingly, both the N- or C-terminal incorporation of a Myc sequence completely impaired the secretion of the HEL-specific scFv, suggesting disturbed protein folding or secretion, which is more likely for the tag fused N-terminally, as the tag sequence directly behind the signal peptide can interfere with translocation into the secretory pathway. Constructs containing a Strep tag II showed drastically improved detection and selection of cells engineered with a CBCR compared to constructs including a Flag tag sequence in the extracellular spacer region (**Fig. 2b, 3e, S4**). In contrast to the Myc containing constructs, the Flag tag still enabled surface expression, suggesting compromised Flag tag performance or accessibility. Extracellular linker sequences are expected to provide certain degrees of scFv flexibility, while still allowing signal transduction. Variations of linker length did not increase tag accessibility as measured by tag detection with two different monoclonal antibody clones (M2 and FG4R) and correlation with antigen binding (**Fig. 3**). In a recent study, similar spacer regions including a Flag tag were used to successfully detect the surface expression of synthetic antigen receptors in HEK293FT cells using the antibody clone M2, suggesting that detection and accessibility of an orthogonal tag sequence are additionally influenced by the cell type^45^. Furthermore, the length and composition of extracellular spacers have been reported to be decisive for surface expression and activity of antigen receptors^11,45^. We tested a series of linker sequences, however and did not observe any effect on surface expression (**Fig. 3**). In contrast, we found that the transmembrane domain affected surface expression implying that the transmembrane region also has the capacity to provide stability to the CBCR (**Fig. 2, 3**). CBCR encoding a CD28-derived transmembrane domain showed increased surface expression compared to the CBCR including an endogenous BCR-TM. This result is consistent with previous research revealing that the CD28 TM domain induces a higher expression of CAR than the CD3ζ TM domain^57^. Our findings support that synthetic receptors require careful evaluation of their various components in order to have an optimized expression and detection system.

The signaling proteins CD79a and CD79b are required for the transport of a BCR to the cell surface and for signal transduction^58,59^. Our results show that the inferior surface expression of constructs containing the IgG2c-TM could be partially rescued by fusing the short intracellular tail to the cytoplasmic domain of a CD79 protein (**Fig. 4, 6**). Previous work suggested that CD79a and CD79b are independently sufficient to drive B cell maturation and activation, as long as the ITAM regions of the intracellular signaling domain remain intact^49^. Additonally, our results reflect the improved surface transport of the CD79b construct compared to the CD79a receptor, in accordance with this previous work (**Fig. 4, 6**).

In order to evaluate highly expressed CBCR variants in primary B cells, we developed a reliable pipeline to genomically integrate large gene cassettes by Cas9-driven HDR (**Fig. 5**). While many years of work have aimed to reprogram immune cells for therapeutic purposes, such as CAR T cell therapy, these have almost exclusively relied on viral-based gene transfer. Recently, genome editing platforms providing targeted integration, most notably CRISPR-Cas9, have become promising tools to further improve current immune cell therapies, by offering potential advantages related to safety, uniform expression levels and potency^56,60,61,62^. Establishing a preclinical genome editing platform based on primary murine B cells does not only show progress on cellular engineering of technically challenging target cell lines, but also allows the investigation of these cells as novel vehicle for adoptive immune cell therapies. We observed robust transfection efficiencies (electroporation by nucleofection) in primary B cells following pre-activation and expansion on fibroblast feeder cells expressing BAFF and CD40 ligand. Cas9-RNP-mediated HDR of double-stranded DNA occurred consistently, but with relatively low efficiencies when compared to other primary lymphocyte cells, such as T cells^61,51^. Furthermore, in contrast to observations in multiple other cell types including T and stem cells, AAV delivery of the HDR donor only marginally increased HDR frequencies in primary B cells, suggesting that low HDR efficiencies are independent of template format and transfection efficiencies^54,60^. Notably, the AAV format caused a relatively high background of gene expression from episomal retention of DNA (**Fig. 5f, g**). Our results imply that, for constructs that use a constitutive promoter for gene expression, AAV-based template delivery in primary murine B cells may not be sensitive enough to effectively distinguish edited cells from episomal expression. However, it may be beneficial for approaches that are designed such that only correct integration leads to gene expression (i.e. splicing or use of endogenous promoter)^29,63^. We found dramatically enhanced HDR-mediated integration efficiencies in hybridoma B cells using the same AAV-DJ template targeting the same genomic locus (**Fig. S3**), thus the low HDR frequencies in primary murine B cells are not related generally to inefficient delivery of genome editing reagents or to lack of or targeting specificity. In the future, it would be valuable to determine the potential causes for these inherent limitations of HDR in primary murine B cells; a sensible hypothesis is an upregulation of inhibitory factors for HDR. In fact, very low activity of conservative HDR, known for its high-fidelity, has been described before, while error-prone, non-conservative homologous recombination causing deletions, gene fusions and other genetic aberrations^64^ seem to predominate. Nevertheless, in the context of human B cells, Cas9-RNP with AAV-6 donor has been reported to be highly efficient (at least 10% HDR rate; 100-fold higher than what we observed in this study), emphasizing once more that the differences between murine and human cells must not be underestimated^19,34^.

The clear discrimination of edited cells using PCR-derived HDR donor still offers a very reliable tool to develop new concepts for cellular therapies. Recently, Hung *et al.* used an interesting strategy in primary human B cells by combining gene disruption for plasma cell differentiation with engineering of these cells to secrete a therapeutic protein, followed by *in vivo* transfer in immunodeficient mice^19^. To further evaluate and optimize *in vivo* stability, our approach for cellular engineering in primary murine B cells enables studies that include adoptive transfer to immunocompetent mouse models, which will be valuable for developing novel B cell-based immunotherapies.

## Methods

### Preparation of HDR donor templates

All primers were ordered from Integrated DNA Technologies (IDT) and sequences are listed in Supplementary Table **S1**. HDR donor constructs were cloned by Gibson assembly using the Gibson Assembly Master Mix (NEB, E2611S) into the pUC57(Kan) cloning vector, obtained from Genewiz. The vector was designed with homology arms PCR-amplified from C57BL/6-Ly5.1 genomic DNA according to the mouse genomic sequence (Gt(ROSA)26Sor gene) and Sanger sequenced (pUC57-Rosa26). Codon-optimized nucleotide sequences encoding each transgene or parts of it were synthesized (gBlocks, IDT) or generated by PCR from previously characterized CBCR expression vectors. Anti-HEL scFv was derived from the high-affinity HyHEL10 antibody in the V_L_-V_H_ format or codon-optimized for mice from the D1.3 variant M3 scFv (CA2787677A1). M3 scFv was linked by extracellular spacer regions incorporating different detection tags (Myc, Flag, Strep II) to either a BCR or CD28 transmembrane domain. These TM domains are fused to the short intracellular tail of the BCR C-terminally followed by the cytosolic domain of either the CD79a or CD79b polyprotein. All CBCRs contain an N-terminal V_κ_ signal peptide for membrane targeting. The GFP reporter gene and T2A-CBCR constructs were cloned into the pUC57-Rosa26 under the control of a CMV promoter. HDR donor vectors were linearized by PCR with the KAPA Hifi HotStart ReadyMix (KAPA Biosystems, KK2602) using either p9 and p10 or p11 and p12 (HA 750bp each) for direct comparison with AAV delivered HDR donor. For each PCR, the reaction was split between a minimum of five separate tubes and then pooled for subsequent steps. This split-pool PCR approach was used to minimize the chance of mutations in the repair template arising from PCR. The PCR product was purified using DNA Clean &Concentrator^TM^-5 (Zymo, D4013), eluted in nuclease-free (NF) H_2_O, and concentrated to ~1 μg μl^-1^ using a Concentrator 5301 (Eppendorf).

### Mice

C57BL/6-Ly5.1 mice were obtained by in-house breeding and were maintained in the mouse facility under specific pathogen-free conditions. Mouse procedures were performed under protocols approved by the Basel-Stadt cantonal veterinary office (Basel-Stadt Kantonales Veterinäramt Tierversuchsbewilligung #2701).

### Cell culture

All cell lines were maintained in incubators at 37 °C and 5% CO_2_ and were confirmed to be negative for *Mycoplasma* contamination using a mycoplasma detection kit (ATCC, 30-1012K). If required, the live cell number was counted by the trypan blue dye exclusion method using the TC 10 Automated Cell Counter (Bio-Rad). All B cell hybridoma lines were cultivated in high-glucose Dulbecco’s Modified Eagle Medium (DMEM) containing GlutaMAX supplemented with 10% FBS, 100 U ml^-1^ penicillin/streptomycin, 10 mM HEPES buffer and 50 μM 2-mercaptoethanol. Hybridomas were typically maintained as 5ml cultures in T-25 flasks and passaged every 48 to 72 hours. A list of all hybridoma cell lines is provided in Supplementary Table **S3**. Balb/c 3T3 fibroblast derived 40LB feeder cells were previously generated by Nojima *et al*.^50^, maintained in high-glucose DMEM containing GlutaMAX supplemented with 10% FBS and 100 U ml^-1^ and passaged at 90% confluence. To prepare the feeder layer, 40LB cells were plated at 4 x 10^4^ per cm^2^ about 16 h before co-culture and irradiated with 60 Gy γ–ray. Splenic B cells were pre-activated in a T-75 flask in the presence of irradiated 40LB feeder cells in 40 ml RPMI-1630 medium supplemented with 10% FBS (consistently coming from the same batch), 1 mM Na-Pyruvate, 10 mM HEPES buffer, 100 U ml^-1^ Penicillin/Streptomycin and 50 μM 2-Mercapotethanol for 24 h. rIL-4 (1 ng ml^-1^, Peprotech) was added to the primary culture for four days. From day 4, cells were cultivated on a new feeder layer with rIL-21 (10 ng ml^-1^, Peprotech).

### Splenic B cell isolation

Single cell suspensions of splenocytes were generated from C57BL/6-Ly5.1 mouse spleen under sterile conditions by passing cells through a 70 μM cell strainer using the plunge of a syringe. Subsequently, cells were counted and pelleted at 300g for 10 min at 4 °C before resuspending them in autoMACS running buffer (Miltenyi Biotech). Highly pure resting B cells were isolated by magnetic labeling and depletion of CD43-expressing B cells and non-B cells using the Mouse B cell Isolation Kit (Miltenyi Biotech, 130-090-862) and MACS LS columns (Miltenyi Biotech) following vendor instructions. For activation, up to 3 x 10^7^ cells were plated in a T-75 flask on a 40LB feeder cell layer for 24 h, if not described differently.

### Gene editing in primary murine B cells

24 h after initiating B cell activation on the feeder layer, B cells including the 40LB feeder cells were harvested by collecting the growth medium and dissociating the adherent cells by adding 3 ml autoMACS running buffer (Miltenyi Biotech) to the T-75 flask. Prior to transfection, customized Alt-R crRNA and Alt-R tracrRNA (Alt-R^®^ CRISPR-Cas9 System, IDT) were complexed at equimolar concentrations by incubation at 95 °C for 5 min. crRNAs were designed using the Broad institute single guide RNA (sgRNA) design tool (http://portals.broadinstitute.org/gpp/public/analysis-tools/sgrna-design). Sequences of all tested gRNAs are listed in Supplementary **Table S2**. All genome editing experiments performed utilized Cas9 from *Streptococcus pyogenes* (SpCas9) purchased from IDT. Pre-activated B cells were transfected using the P4 Primary Cell 4D-Nucleofector X Kit L (Lonza, V4XP-4024) in combination with the program DI-100. The following standard conditions in 100 μl total volume of nucleofection mix were used, if not described differently: 1 x 10^6^ cells, 20 μg Cas9 protein complexed with 0.156 nmol Alt-R duplex gRNA at 1:1.125 ratio and 5 μg of linearized double-stranded DNA generated by PCR. After electroporation, edited cells were seeded in 5-6ml culturing medium supplemented with rIL-4 into a 6-well plate in the presence of irradiated 40LB cells (5 x 10^5^ cells per well). Two days after transfection (Day 3), the B cell culturing medium was replaced by carefully aspirating the medium and adding 5 ml of fresh B cell medium supplemented with rIL-4. One day later (Day 4), primary B cells were harvested and prepared for flow cytometry analysis.

### Gene editing in hybridoma cells

Genome editing experiments in B cell hybridomas were always executed in the HC9-cell line being dysfunctional in antibody expression and constitutively expressing Cas9 protein^30^. Hybridoma cells were electroporated using the SF Cell Line 4D-Nucleofector X Kit L (Lonza, V4XC-2024) with the program CQ-104. The following standard conditions in 100 μl of total volume of nucleofection mix were used: 1 x 10^6^ cells, 0.156 nmol pre-complexed Alt-R duplex gRNA and 5 μg of PCR-linearized double-stranded DNA. Following transfection, cells were incubated for 5 min at RT, before adding 500 μl of pre-warmed medium to the nucleocuvette and transferring them to 1.5 ml of fresh growth medium in 6-well plates. The cells were usually supplemented 24 h later with 0.5-1.0 ml of fresh culturing medium.

### Transduction with AAV

AAV vector plasmids were cloned in the pMD13-AAV plasmid containing inverted terminal repeats from AAV serotype 2. HDR donor cassettes including a GFP gene under the control of a CMV promoter and a SV40 polyA sequence flanked by 750bp homology arms of the Rosa26 locus were inserted into the multiple cloning site (MCS) by NotI restriction digest. Cloning was performed in the suitable bacterial strain Stbl3. AAV stocks were produced by the Viral Vector Facility (VVF) of the Neuroscience Center Zurich. The optimal AAV serotype for the transduction of primary murine B cells was evaluated by adding ssAAV with a GFP coding sequence packaged using various serotypes (AAV1, 2, 6, 7, 8, 9, DJ) at an MOI of 2.5 x 10^10^ to B cells pre-activated for 24 h. For samples transduced with AAV for HDR template delivery, AAV-DJ donor vector was added to the culture immediately after electroporation at an MOI (vector genomes/cell) of 20,000 – 5 x 10^5^ and cultured as described for transfected cells. AAV donor was added as 10% of the final culture volume regardless of titer.

### Genomic analysis of CRISPR-Cas9 targeting

The activity of gRNAs targeting the Rosa26 locus were initially tested by induction of NHEJ. Cells transfected with Cas9-RNP targeting the Rosa26 locus were harvested four days after electroporation, washed once in PBS and genomic DNA was recovered from 1 x 10^6^ cells using 100 μl Quick Extract solution (Epicentre) according to the manufacturer’s instructions. Small fragments of DNA covering the putative cleavage sites were amplified by PCR with KAPA Hifi HotStart Ready Mix (KAPA Biosystems, KK2602) from the genomic DNA using primers p13 and p14. Control DNA was also amplified from wildtype C57BL/6-Ly5.1 genomic DNA. CRISPR-Cas9 cleavage of the genome was determined using a Surveyor Mutation Detection Kit (IDT, 706020). All samples were run on 2% gels for the detection of cleavage products. For reference, GeneRuler 1 kb DNA Ladder (Thermo, SM0314) and GeneRuler 100bp DNA Ladder (Thermo, SM0243) were used as DNA size markers.

### Measuring targeted integration of CBCR construct

For the evaluation of transgene integration, PCR analysis was performed on genomic DNA extracted from sorted cells or single-cell clones excluding the presence of remaining repair template. Primer p3 and p4, closely flanking the gRNA targeting site in the Rosa26 locus in combination with KAPA Hifi HotStart Ready Mix were used with the following PCR conditions: 35 cycles with annealing at 62 °C (15 s), elongation at 72 °C (1:30 min) and final elongation at 72 °C (3:00 min).

To determine targeted integration mediated via HDR, PCR was performed on genomic DNA using primers binding inside the construct cassette and outside of homology arm. Primer p5 and p6 were used with the following cycling conditions: 35 cycles with annealing at 69 °C (15 s), elongation at 72 °C (1:30 min), final elongation at 72 °C (3:00 min), primer p7 and p8: 35 cycles with annealing at 71 °C (15 s), elongation at 72 °C (1:30 min), final elongation at 72 °C (3:00 min), primer p7 and p15: 35 cycles with annealing at 73 °C (15 s), elongation at 72 °C (1:30 min), final elongation at 72 °C (3:00 min).

### Evaluation of scFv expression by RT-PCR

To confirm transcript expression of the HEL-specific scFv variants, mRNA was isolated from 1 x 10^6^ transfected or GFP-bulk sorted hybridoma and parental HC9-cells using 200μl TRIzol® reagent (Thermo, 15596-026). The mRNA was purified using the PureLink Mini Kit (Invitrogen, Thermo) according to the manufacturer’s instructions. First-strand cDNA was synthesized from mRNA using Maxima Reverse Transcriptase (Thermo, EP0742) and used as template DNA for subsequent PCR reactions. For the detection of correct transcript expression, the following cycling conditions were applied using KAPA Hifi HotStart Ready Mix and p1 and p2, binding to GFP and the SV40 polyA sequence: 25 cycles with annealing at 61 °C (15s), elongation at 72 °C (1 min), final elongation at 72 °C (2 min).

### Measuring scFv secretion by ELISA

Three days prior to measuring culture scFv levels, GFP^+^ sorted cells were collected, counted and then resuspended in new culture medium. After three days, the cell culture supernatant was collected from 1 x 10^6^ cells and normalized to least-concentrated sample. scFv secretion levels were analyzed by ELISA after coating with HEL antigen (Sigma-Aldrich, 62971, 4 μg ml^-1^) in PBS (Thermo, 10010-015). The plates were then blocked with PBS supplemented with 2% m/v milk (AppliChem, A0830) and 0.05% V/V Tween-20 (AppliChem, A1389, PBSMT) followed by three washing steps with PBS supplemented with Tween-20 0.05% V/V (PBST). Supernatants were then serially diluted (at 1:3 ratio) in PBSMT, starting from the non-diluted supernatant as the highest concentration. Supernatants were incubated for 1 h at RT, followed by three washing steps with PBST and incubation with HRP-conjugated anti-Myc antibody (9E10, Thermo Fisher Scientific, MA1-81357) or anti-Flag antibody (FG4R, Thermo Fisher Scientific, MA1-91878-HRP) at 2 μg ml^-1^ (1:500 dilution from stock) in PBSTM. After three more washing steps with PBST, ELISA detection was performed using a 1-Step Ultra TMB-ELISA Substrate Solution (Thermo, 34028), reaction was terminated with H_2_SO_4_ (1 M). Absorbance at 450 nm was measured using an Infinite 200PRO NanoQuant plate reader (Tecan). ELISA data were analysed with the software GraphPad Prism.

### Flow cytometry analysis and sorting for immunophenotyping

Flow cytometry-based analysis and cell isolation were performed on a 5 laser BD LSR Fortessa™ flow cytometer and BD FACS Aria III (BD Biosciences), respectively. Data were analyzed with FlowJo software (Tree Star).

24 h post transfection in any of the cell lines, ~100 μl were collected and analyzed for GFP expression (via GFP-T2A). Primary B cells were only harvested for sorting on GFP expression three days after transfection. Hybridoma cells were enriched for GFP expressing cells three days post transfection, if not indicated differently. After sorting and expansion primary B cells or hybridoma cells were labeled with HEL-antigen, conjugated to Alexa Fluor 647 dye using the Alexa Fluor®647 Protein Labeling Kit (Thermo Fisher Scientific, A20173) according to the manufacurer’s instructions, and antibodies binding the respective detection tag to determine CBCR surface expression. For this purpose, cells were washed with phosphate-buffered saline (PBS), incubated with the labeling antibody or antigen for 30 min on ice or 10 min at RT, protected from light, washed again with PBS and analyzed or sorted. Biotinylated antibodies were stained with Strepatvidin-BV421 (Biolegend). Staining with propidium iodide (PI, BD BioSciences) was used for live/dead cell discrimination as directed by the manufacturer. When primary B cells that had been cultured on a 40LB feeder layer were analyzed, 40LB feeder cells were excluded based on FSC versus SSC. The labeling reagents and working concentrations are described in Supplementary Table S4.

## Acknowledgements

We acknowledge the ETH Zurich D-BSSE Single Cell Unit and the ETH Zurich D-BSSE Animal Facility for support, in particular, T. Lopes, V. Jäggin, Marie-Didiée-Hussherr and Gieri Camenisch. We are grateful to Mark Pogson for providing initial scientific discussions and feedback. Funding was provided by the National Competence Center for Research on Molecular Systems Engineering.

## Author Contributions (Articles only)

T.P. and S.T.R. developed methodology and designed experiments; T.P. and L.B. performed experiments; W.K., C.P., R.E. and L.C. generated critical materials and provided technical advice. T.P, L.B and S.T.R. analyzed data and wrote manuscript.

