## Supplementary material for "Molecular design, optimization and genomic integration of chimeric B cell receptors in murine B cells" (from another version of this preprint: version 1, 2019-01-17)

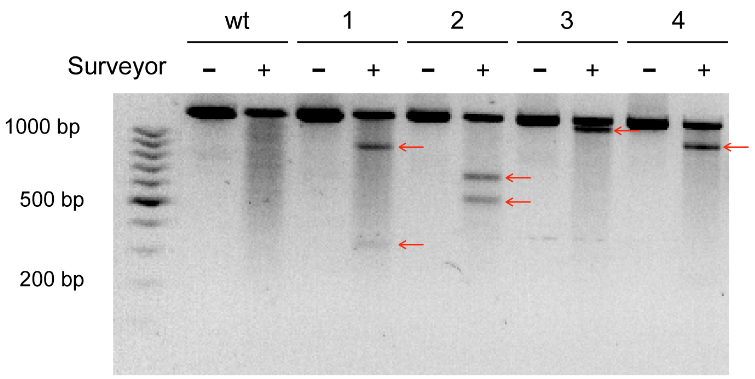


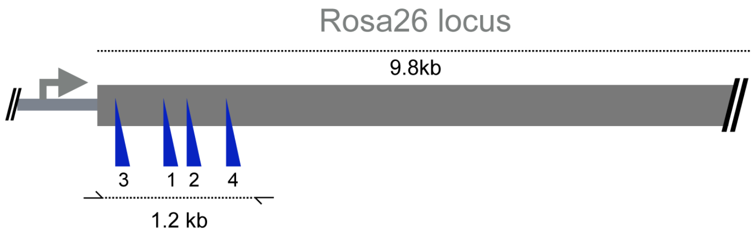


| gRNA | **Frag 1** in bp | **Frag 2** in bp |
|---|---|---|
| 1 | 875 | 325 |
| 2 | 670 | 530 |
| 3 | 100 | 1100 |
| 4 | 950 | 250 |

**Figure S1. Validation of CRISPR-Cas9 targeting in Rosa26 locus.** Surveyor assays following transfection of Cas9 protein complexed with gRNAs targeting the safe harbor Rosa26 locus. Non-transfected cells were used as negative control (wt). Table shows the approximate length of expected fragments after cleavage. Red arrows in agarose gel indicate cleavage products. gRNA1 was selected for further experiments.

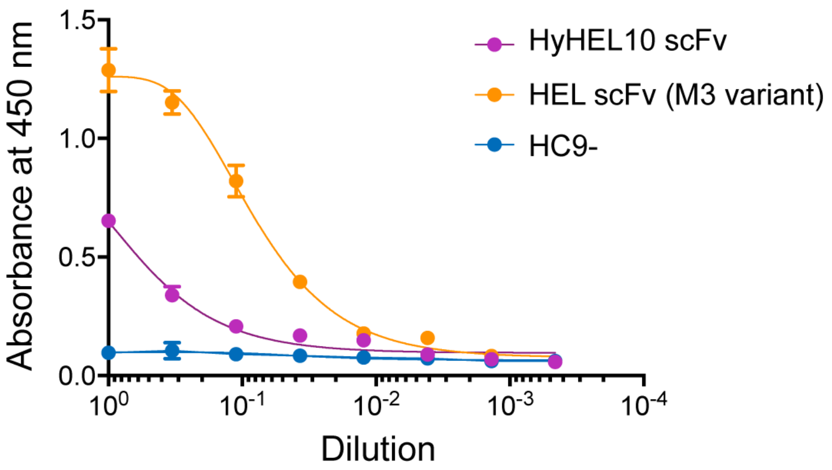


**Figure S2. Secretion test of HEL-binding scFv variants.** Cells were enriched for GFP (488 nm) expression via FACS. Graph shows ELISA results of scFv secretion levels (capture HEL antigen, detection anti-Flag) on enriched hybridoma culture supernatant for scFv variants, M3 variant of D1.3 ($V_H$-$V_L$ format) and HyHEL10 variant ($V_L$-$V_H$). Supernatant of HC9- cells was used as negative control. For each sample, three technical replicates were analysed and a four-parameter logistical curve was fitted to the data by nonlinear regression. Data are presented as the mean and error bars indicate standard deviation.

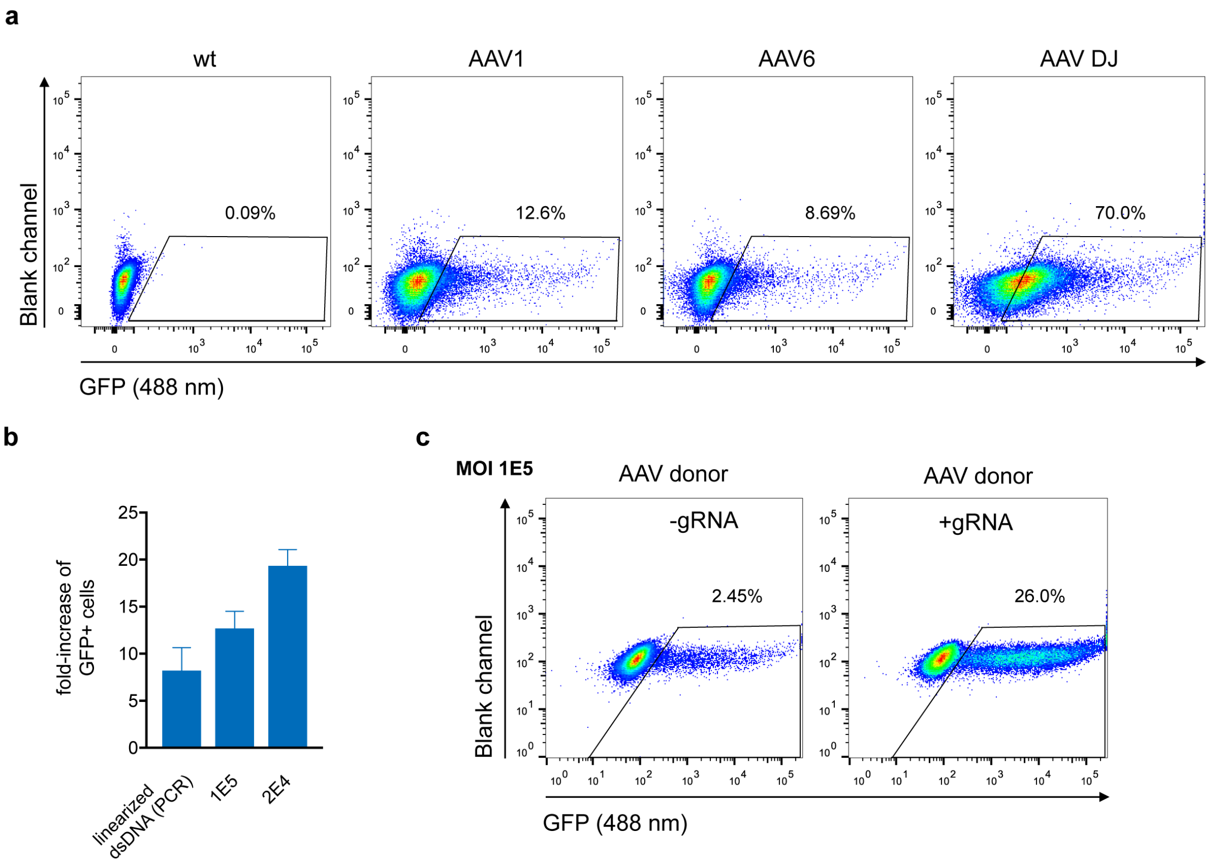


**Figure S3. Delivery of HDR donor DNA in the format of AAV. a**) Representative flow cytometry plots show the transduction efficiencies for serotype AAV-1, AAV-6 and AAV-DJ at a MOI of $10^5$ three days after transduction or mock treatment. **b**) HC9- cells were transfected with Cas9-RNP immediately followed by HDR repair template delivery via chimeric AAV serotype DJ encoding the GFP reporter gene. Cells transfected with Cas9-RNP and PCR-linearized dsDNA served as control. GFP expression was measured on day 9 after transfection. Cells transfected with Cas9-protein only indicate the episomal AAV background expression. Data are displayed as fold-increase of $GFP^+$ cells receiving the Cas9-gRNA complex to cells transfected with Cas9-protein only representing the HDR based integration. All data are means ±s.d (n=3). **c**) Representative flow cytometry dot plots show GFP expression (488 nm) day for transduction with a MOI of $10^5$. Cells transfected only with Cas9-protein without gRNA and transduced with GFP expressing HDR donor packaged using scAAV-DJ were used as negative control to determine the background level of episomal GFP expression.

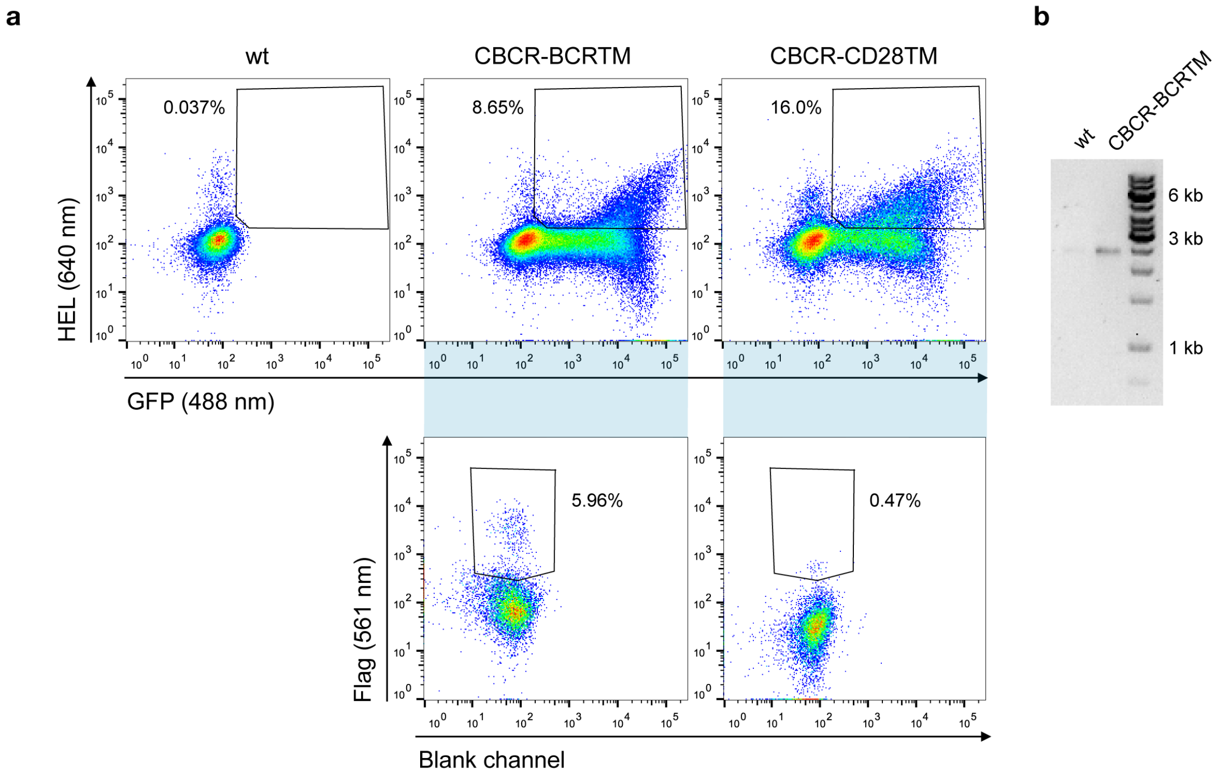


**Figure S4. Robust CBCR surface expression and antigen binding, but modest Flag detection in primary B cells. a**) Three days after transfection of splenic B cells with Cas9-RNP and repair templates encoding CBCR constructs with both TM domains and Flag detection tag within the extracellular spacer, GFP expressing cells (488 nm) were enriched by FACS. Flow cytometry dot plots show efficient enrichment of GFP+ cells and HEL antigen binding (upper row), but only modest detection of the Flag epitope (561 nm) within this population (lower row). **b**) Agarose gel shows PCR product from genomic DNA analysis (p7/15) that confirm the targeted integration of the GFP-T2A-CBCR-BCRTM into the Rosa26 locus (2442bp).

### Table S1: Primers used for DNA construct engineering and genomic analysis

| Name | Binding region | Sequence (5′ to 3′) |
|---|---|---|
| p1 (for) | GFP | ATGGAGATCGAGTGCCGCATCA |
| p2 (rev) | polyA | CACTGCGATTAAAGACATGCTCACCCG |
| p3 (for) | 5′ HA (Rosa26) | CAGATTTTCGGTTTTGTCGGG |
| p4 (rev) | 3′ HA (Rosa26) | GGATATGAAGTACTGGGCTCTT |
| p5 (for) | M3 scFV ($V_H$) | ATCTGGGGAGACGGCAATAC |
| p6 (rev) | Rosa26 | AGTAATGGCTGCAGACTTAGCTTT |
| p7 (for) | Rosa26 | CCACCGCCCCACACTTATT |
| p8 (rev) | M3 scFv ($V_H$) | ACCCAGTCAAGCTAAACCCG |
| p9 (for) | Rosa26 (5′ HA long) | CAGTTATGGCTCCTCTGTCCAC |
| p10(rev) | Rosa26 (3′ HA long) | CAGAGAGCCTCGGCTAGGT |
| p11(for) | Rosa26 (5′ HA AAV) | ATTATGTTAGCGGCCGCATTGGCTCGTGTTCGTGCA |
| p12(rev) | Rosa26 (3′ HA AAV) | ACTTCAGCGGCCGCCACCTTTTACAGATGTGTAC |
| p13 (for) | Rosa26 (Surveyor) | GTTCTCTGCTGCCTCCTG |
| p14 (rev) | Rosa26 (Surveyor) | CAAGCCAGTCCAAGAGAAAGCACTG |
| p15 (rev) | T2A | TTGAAGTCGCCGATCACG |

### Table S2: List of all gRNAs used in this study

| Name | Target region | Sequence (5′ to 3′) |
|---|---|---|
| 1 | Rosa26 | CGATGGAAAATACTCCGAGG |
| 2 | Rosa26 | GTTACATACACCACAAATCG |
| 3 | Rosa26 | ACTCCAGTCTTTCTAGAAGA |
| 4 | Rosa26 | AGATACATCAGGTAATATTG |

### Table S3: Summary of cell lines used and generated in this study

| Name | Description |
|---|---|
| Wen1.3 | Wen1.3 cells are derived from a mouse infected with LCMV. They express IgG2c and are specific for LCMV GP-1 antigen. |
| HC9- | HC9- are a monoclonal population and contain a frameshift mutation in the CDRH3 resulting in dysfunctional antibody expression (PnP-HEL23.Fl[71]). It also includes constitutive Cas9 expression. |
| CBCR-CD79b | Representative example (including the CD79b intracellular domain) for all cell lines containing a CBCR construct precisely integrated into the Rosa26 locus. CBCR constructs are constitutively expressed under the control of a CMV promoter. CBCR expressing cells are derived from HC9-. |
| 40LB | 40LB are Balb/c 3T3 fibroblasts stably expressing CD40L and BAFF and used for *in vitro* culture of primary murine B cells[112]. |

### Table S4: Flow cytometry labeling reagents with their working concentrations.

| Target antigen | Conjugate | Working concentration (dilution from stock) | Cat. No, supplier |
|---|---|---|---|
| Flag (FG4R) | DyLight550 | 5 µg $ml^{-1}$ (1:200) | MA1-91878-D550, Thermo |
| Strep tag II | Biotin | 2.5 µg $ml^{-1}$ (1:200) | A01732, GenScript |
| Hen egg lysozyme (HEL) | AlexaFluor®647 | 1.6 µg $ml^{-1}$ (1:200) | 62971-10G-F (Sigma Aldrich) |
| Biotin | Brilliant Violet 421 | 0.2 µg $ml^{-1}$ (1:500) | 405226, BioLegend |

### Table S5: Antibodies used for ELISA assays

| Target antigen | Conjugate | Host | Working concentration (dilution from stock) | Cat. No, supplier |
|---|---|---|---|---|
| Myc tag (9E10) | HRP | mouse | 2 µg $ml^{-1}$ (1:500) | MA1-81357, Thermo |
| Flag (FG4R) | HRP | mouse | 2 µg $ml^{-1}$ (1:500) | MA1-91878-HRP, Thermo |

### Protein and DNA sequences

#### Rosa26 locus

CCACCGCCCCACACTTATTGGCCGGTGCGCCGCCAATCAGCGGAGGCTGCCGGGGCCGCCTA
AAGAAGAGGCTGTGCTTTGGGGCTCCGGCTCCTCAGAGAGCCTCGGCTAGGTAGGGGATCGGG
ACTCTGGCGGGAGGGCGGCTTGGTGCGTTTGCGGGGATGGGCGGCCGCGGCAGGCCCTCCGA
GCGTGGTGGAGCCGTTCTGTGAGACAGCCGGGTACGAGTCGTGACGCTGGAAGGGGCAAGCG
GGTGGTGGGCAGGAATGCGGTCCGCCCTGCAGCAACCGGAGGGGGAGGGAGAAGGGAGCGG
AAAAGTCTCCACCGGACGCGGCCATGGCTCGGGGGGGGGGGGGCAGCGGAGGAGCGCTTCC
GGCCGACGTCTCGTCGCTGATTGGCTTCTTTTCCTCCCGCCGTGTGTGAAAACACAAATGGCGT
GTTTTGGTTGGCGTAAGGCGCCTGTCAGTTAACGGCAGCCGGAGTGCGCAGCCGCCGGCAGCC
TCGCTCTGCCCACTGGGTGGGGCGGGAGGTAGGTGGGGTGAGGCGAGCTGGACGTGCGGGC
GCGGTCGGCCTCTGGCGGGGCGGGGGAGGGGAGGGAGGGTCAGCGAAAGTAGCTCGCGCGC
GAGCGGCCGCCCACCCTCCCCTTCCTCTGGGGGAGTCGTTTTACCCGCCGCCGGCCGGGCCT
CGTCGTCTGATTGGCTCTCGGGGCCCAGAAAACTGGCCCTTGCCATTGGCTCGTGTTCGTGCAA
GTTGAGTCCATCCGCCGGCCAGCGGGGGCGGCGAGGAGGCGCTCCCAGGTTCCGGCCCTCCC
CTCGGCCCCGCGCCGCAGAGTCTGGCCGCGCGCCCCTGCGCAACGTGGCAGGAAGCGCGCG
CTGGGGGCGGGGACGGGCAGTAGGGCTGAGCGGCTGCGGGGCGGGTGCAAGCACGTTTCCG
ACTTGAGTTGCCTCAAGAGGGGCGTGCTGAGCCAGACCTCCATCGCGCACTCCGGGGAGTGGA
GGGAAGGAGCGAGGGCTCAGTTGGGCTGTTTTGGAGGCAGGAAGCACTTGCTCTCCCAAAGTC
GCTCTGAGTTGTTATCAGTAAGGGAGCTGCAGTGGAGTAGGCGGGGAGAAGGCCGCACCCTTC
TCCGGAGGGGGGAGGGGAGTGTTGCAATACCTTTCTGGGAGTTCTCTGCTGCCTCCTGGCTTCT
GAGGACCGCCCTGGGCCTGGGAGAATCCCTTCCCCCTCTTCCCTCGTGATCTGCAACTCCAGTC
TTTCTAGAAGATGGGCGGGAGTCTTCTGGGCAGGCTTAAAGGCTAACCTGGTGTGTGGGCGTTG
TCCTGCAGGGGAATTGAACAGGTGTAAAATTGGAGGGACAAGACTTCCCACAGATTTTCGGTTTT
GTCGGGAAGTTTTTTAATAGGGGCAAATAAGGAAAATGGGAGGATAGGTAGTCATCTGGGGTTTT
ATGCAGCAAAACTACAGGTTATTATTGCTTGTGATCCGCCTCGGAGTATTTTCCATCGAGGTAGA
TTAAAGACATGCTCACCCGAGTTTTATACTCTCCTGCTTGAGATCCTTACTACAGTATGAAATTAC
AGTGTCGCGAGTTAGACTATGTAAGCAGAATTTTAATCATTTTTAAAGAGCCCAGTACTTCATATC
CATTTCTCCCGCTCCTTCTGCAGCCTTATCAAAAGGTATTTTAGAACACTCATTTTAGCCCCATTT
TCATTTATTATACTGGCTTATCCAACCCCTAGACAGAGCATTGGCATTTTCCCTTTCCTGATCTTA
GAAGTCTGATGACTCATGAAACCAGACAGATTAGTTACATACACCACAAATCGAGGCTGTAGCTG
GGGCCTCAACACTGCAGTTCTTTTATAACTCCTTAGTACACTTTTTGTTGATCCTTTGCCTTGATC
CTTAATTTTCAGTGTCTATCACCTCTCCCGTCAGGTGGTGTTCCACATTTGGGCCTATTCTCAGTC
CAGGGAGTTTTACAACAATAGATGTATTGAGAATCCAACCTAAAGCTTAACTTTCCACTCCCATGA
ATGCCTCTCTCCTTTTTCTCCATTTATAAACTGAGCTATTAACCATTAATGGTTTCCAGGTGGATG
TCTCCTCCCCCAATATTACCTGATGTATCTTACATATTGCCAGGCTGATATTTTAAGACATTAAAA
GGTATATTTCATTATTGAGCCACATGGTATTGATTACTGCTTACTAAAATTTTGTCATTGTACACAT
CTGTAAAAGGTGGTTCCTTTTGGAATGCAAAGTTCAGGTGTTTGTTGTCTTTCCTGACCTAAGGT
CTTGTGAGCTTGTATTTTTTCTATTTAAGCAGTGCTTTCTCTTGGACTGGCTTGACTCATGGCATT
CTACACGTTATTGCTGGTCTAAATGTGATTTTGCCAAGCTTCTTCAGGACCTATAATTTTGCTTGA
CTTGTAGCCAAACACAAGTAAAATGATTAAGCAACAAATGTATTTGTGAAGCTTGGTTTTTAGGTT
GTTGTGTTGTGTGTGCTTGTGCTCTATAATAATACTATCCAGGGGCTGGAGAGGTGGCTCGGAG
TTCAAGAGCACAGACTGCTCTTCCAGAAGTCCTGAGTTCAATTCCCAGCAACCACATGGTGGCTC
ACAACCATCTGTAATGGGATCTGATGCCCTCTTCTGGTGTGTCTGAAGACCACAAGTGTATTCAC
ATTAAATAAATAAATCCTCCTTCTTCTTCTTTTTTTTTTTTTTAAAGAGAATACTGTCTCCAGTAGAA
TTTACTGAAGTAATGAAATACTTTGTGTTTGTTCCAATATGGTAGCCAATAATCAAATTACTCTTTA
AGCACTGGAAATGTTACCAAGGAACTAATTTTTATTTGAAGTGTAACTGTGGACAGAGGAGCCAT
AACTGCAGACTTGTGGGATACAGAAGACCAATGCAGACTTTAATGTCTTTTCTCTTACACTAAGCA
ATAAAGAAATAAAAATTGAACTTCTAGTATCCTATTTGTTTAAACTGCTAGCTTTACTTAACTTTTGT
GCTTCATCTATACAAAGCTGAAAGCTAAGTCTGCAGCCATTACT

Grey: Homology arms

Orange: PAM

Blue: CRISPR gRNA target site

Red: primer for genomic DNA analysis (p9 and p10)

**CMV promoter**
ATAGTAATCAATTACGGGGTCATTAGTTCATAGCCCATATATGGAGTTCCGCGTTACATAACTTAC
GGTAAATGGCCCGCCTGGCTGACCGCCCAACGACCCCCGCCCATTGACGTCAATAATGACGTAT
GTTCCCATAGTAACGCCAATAGGGACTTTCCATTGACGTCAATGGGTGGAGTATTTACGGTAAAC
TGCCCACTTGGCAGTACATCAAGTGTATCATATGCCAAGTCCGCCCCCTATTGACGTCAATGACG
GTAAATGGCCCGCCTGGCATTATGCCCAGTACATGACCTTACGGGACTTTCCTACTTGGCAGTA
CATCTACGTATTAGTCATCGCTATTACCATGGTGATGCGGTTTTGGCAGTACACCAATGGGCGTG
GATAGCGGTTTGACTCACGGGGATTTCCAAGTCTCCACCCCATTGACGTCAATGGGAGTTTGTTT
TGGCACCAAAATCAACGGGACTTTCCAAAATGTCGTAATAACCCCGCCCCGTTGACGCAAATGG
GCGGTAGGCGTGTACGGTGGGAGGTCTATATAAGCAGAGGTCGTTTAGTGAACCGTCAGATC

**GFP**
ATGGAGAGCGACGAGAGCGGCCTGCCCGCCATGGAGATCGAGTGCCGCATCACCGGCACCCT
GAACGGCGTGGAGTTCGAGCTGGTGGGCGGCGGAGAGGGCACCCCCAAGCAGGGCCGCATGA
CCAACAAGATGAAGAGCACCAAAGGCGCCCTGACCTTCAGCCCCTACCTGCTGAGCCACGTGAT
GGGCTACGGCTTCTACCACTTCGGCACCTACCCCAGCGGCTACGAGAACCCCTTCCTGCACGC
CATCAACAACGGCGGCTACACCAACACCCGCATCGAGAAGTACGAGGACGGCGGCGTGCTGCA
CGTGAGCTTCAGCTACCGCTACGAGGCCGGCCGCGTGATCGGCGACTTCAAGGTGGTGGGCAC
CGGCTTCCCCGAGGACAGCGTGATCTTCACCGACAAGATCATCCGCAGCAACGCCACCGTGGA
GCACCTGCACCCCATGGGCGATAACGTGCTGGTGGGCAGCTTCGCCCGCACCTTCAGCCTGCG
CGACGGCGGCTACTACAGCTTCGTGGTGGACAGCCACATGCACTTCAAGAGCGCCATCCACCC
CAGCATCCTGCAGAACGGGGGCCCCATGTTCGCCTTCCGCCGCGTGGAGGAGCTGCACAGCAA
CACCGAGCTGGGCATCGTGGAGTACCAGCACGCCTTCAAGACCCCCATCGCCTTCGCCAGATC
CCGCGCTCAGTCGTCCAATTCTGCCGTGGACGGCACCGCCGGACCCGGCTCCACCGGATCTCG
C

MESDESGLPAMEIECRITGTLNGVEFELVGGGEGTPKQGRMTNKMKSTKGALTFSPYLLSHVMGYG
FYHFGTYPSGYENPFLHAINNGGYTNTRIEKYEDGGVLHVSFSYRYEAGRVIGDFKVVGTGFPEDSVI
FTDKIIRSNATVEHLHPMGDNVLVGSFARTFSLRDGGYYSFVVDSHMHFKSAIHPSILQNGGPMFAFR
RVEELHSNTELGIVEYQHAFKTPIAFARSRAQSSNSAVDGTAGPGSTGSR

**T2A**
GGCAGTGGAGAGGGCAGAGGAAGTCTGCTAACATGCGGTGACGTCGAGGAGAATCCTGGCCCA

GSGEGRGSLLTCGDVEENPGP

**Signal peptide**
ATGGTTTTCACACCTCAGATACTTGGACTTATGCTTTTTTGGATTTCAGCCTCCAGAGGT

MVFTPQILGLMLFWISASRG

**HEL scFv (HyHEL10 derived)**
GATATTGTGCTAACTCAGTCTCCAGCCACCCTGTCTGTGACTCCAGGAAATAGCGTCAGTCTTTC
CTGCAGGGCCAGCCAAAGTATTGGCAACAACCTACACTGGTATCAACAAAAATCACATGAGTCTC
CAAGGCTTCTCATCAAGTATGCTTCCCAGTCCATCTCTGGCATCCCCTCCAGGTTCAGTGGCAGT
GGATCAGGGACAGATTTCACTCTCAGTATCAACAGTGTGGAGACTGAAGATTTTGGAATGTATTT
CTGTCAACAGAGTAACAGCTGGCCGTACACGTTCGGAGGGGGGACCAAGCTGGAAATAAAAGG
TGGTGGCGGTTCTGGCGGCGGTGGCTCCGGTGGTGGCGGCAGCGGTGGTGGCGGTAGCGAC
GTGCAGCTTCAGGAGTCAGGACCTAGCCTCGTGAAACCTTCTCAGACTCTGTCCCTCACCTGTT
CTGTCACTGGCGACTCCATCACCAGTGATTATTGGAGCTGGATCAGGAAATTCCCAGGGAATAG
ACTTGAGTACATGGGGTACGTAAGCTACAGTGGTAGCACGTACTACAATCCATCTCTCAAAAGTC
GAATCTCCATCACTCGAGACACATCCAAGAATCAGTACTATCTGGACCTGAATTCTGTGACTACT
GAGGACACAGCCACATATTACTGTGCAAACTGGGACGGTGATTACTGGGGCCAAGGGACTCTG
GTCACTGTCTCTGCA

DIVLTQSPATLSVTPGNSVSLSCRASQSIGNNLHWYQQKSHESPRLLIKYASQSISGIPSRFSGSGSG
TDFTLSINSVETEDFGMYFCQQSNSWPYTFGGGTKLEIKGGGGSGGGGSGGGGSGGGGSDVQLQ

ESGPSLVKPSQTLSLTCSVTGDSITSDYWSWIRKFPGNRLEYMGYVSYSGSTYYNPSLKSRISITRDT
SKNQYYLDLNSVTTEDTATYYCANWDGDYWGQGTLVTVSA

Blue: $V_L$
Grey: GGGGS linker
Orange: $V_H$

**HEL scFv (D1.3 variant M3 derived)**
CAGGTACAATTGCAAGAATCAGGACCAGGATTGGTTGCTCCCTCACAGTCCCTGTCCATCACAT
GCACCGTGAGCGGGTTTAGCTTGACTGGGTATGGTGTCAATTGGGTTAGACAACTGCCTGGCAA
GGGACTCGAGTGGCTCGGCATGATCTGGGGAGACGGCAATACAGCATATAACTCAGCATTGAAA
AGTCGGTTGAGCATATCTAAGGACAACAGCAAATCCCAAGTATTTCTTAAGATGGATTCTTTGCA
CACTGACGACACTGCACGCTACTACTGCGCTCGGGAACGGGATTACAGATTGGATTATTGGGGA
CAGGGAACTACTGTAACCGTTAGCTCTGGTGGCGGAGGGTCAGGAGGTGGTGGTTCAGGTGGT
GGCGGGAGTGACATCAAATTGACCCAATCTCCAGCAAGTTTGAGCGCTTCAGTAGGAGAAACAG
TTACAATCACCTGTAGAGCCTCCGGGAATACTCACAATTACCTTGCCTGGTATCAACAGAAACAA
GGAAAGTCCCCACAGCTCTTGGTATATTACACTACAACACTTGCTGACGGAGTTCCCTCAAGGTT
TAGCGGTTCCGGGAGCGGAACACAGTATTCACTCAAGATCAACTCACTGCAACCAGAAGATTTT
GGAAGCTACTACTGTCAACACTTCTGGTCTACCCCTCGCTCCTTTGGGGGTGGCACCAAATTGG
AAATCAAACGG

QVQLQESGPGLVAPSQSLSITCTVSGFSLTGYGVNWVRQLPGKGLEWLGMIWGDGNTAYNSALKS
RLSISKDNSKSQVFLKMDSLHTDDTARYYCARERDYRLDYWGQGTTVTVSSGGGGSGGGGSGGG
GSDIKLTQSPASLSASVGETVTITCRASGNTHNYLAWYQQKQGKSPQLLVYYTTTLADGVPSRFSGS
GSGTQYSLKINSLQPEDFGSYYCQHFWSTPRSFGGGTKLEIKR

**Extracellular spacer**

ECL23
GGCGCCGGATCTGGAGGAGACTACAAGGACGATGACGACAACGGCGGTTCGGGTGGATCGGG
GGGAGGT

GAGSGGDYKDDDDNGGSGGSGGG

ECL26

GGCGCCGGATCTGGAGGAGACTACAAGGACGATGACGACAACGGTGGTAGTGGCGGTTCGGG
TGGATCGGGGGGAGGT

GAGSGGDYKDDDDNGGSGGSGGSGGG

ECL32

GGCGCCGGATCTGGAGGAGACTACAAGGACGATGACGACAACGGCGGATCGGGGGGGTCTGG
TGGTAGTGGCGGTTCGGGTGGATCGGGGGGAGGT

GAGSGGDYKDDDDNGGSGGSGGSGGSGGSGGG

Linker with Strep tag II (1x)

GGCGCCGGATCTGGAGGAAACTGGTCACACCCACAGTTCGAGAAGGGTGGTAGTGGCGGTTCG
GGTGGATCGGGGGGAGGT

GAGSGGNWSHPQFEKGGSGGSGGSGGG

Linker with Strep tag II (3x)

GGCGCCGGATCTGGAGGAAACTGGTCACACCCACAGTTCGAGAAGGGTGGTAGTGGCGGTTCG
GGTGGAAGTGGAGGTTCGAACTGGTCGCACCCACAGTTCGAGAAGGGGGGGAGCGGGGGCTC
AGGCGGTAGTGGGGGCTCTAACTGGTCCCACCCACAGTTCGAGAAGGGTGGATCGGGGGGAG
GT

GAGSGGNWSHPQFEKGGSGGSGGSGGSNWSHPQFEKGGSGGSGGSGGSNWSHPQFEKGGSG
GG

Orange: Flag tag

Red: Strep tag II

### BCR (IgG2c)-TM

CTAGACCTGGATGATATCTGTGCTGAGGCCAAGGATGGGGAGCTGGACGGGCTCTGGACGACC
ATCACCATCTTCATCAGCCTCTTCCTGCTCAGCGTGTGCTACAGCGCCTCTGTCACACTCTTCAA
GGTA

LDLDDICAEAKDGELDGLWTTITIFISLFLLSVCYSASVTLFKV

## CD28-TM

TTCTGGGCCCTGGTGGTGGTGGCCGGCGTGCTGTTCTGTTACGGCCTGCTGGTCACAGTGGCC
CTGTGCGTGATCTGGACC

FWALVVVAGVLFCYGLLVTVALCVIWT

### BCR-IC

AAGTGGATCTTCTCCTCAGTGGTGGAGCTGAAGCAGAAGATCTCCCCTGACTACAGAAACATGA
TTGGGCAGGGAGCC

KWIFSSVVELKQKISPDYRNMIGQGA

## CD79α-IC

AGGAAGAGGTGGCAGAACGAAAAATTTGGGGTAGACATGCCAGATGACTACGAGGATGAGAATC
TGTATGAAGGTCTCAATCTTGACGATTGCTCAATGTACGAAGACATATCCAGAGGGCTCCAGGG
CACTTACCAGGATGTCGGCAATCTGCATATAGGAGATGCCCAGCTTGAGAAGCCC

RKRWQNEKFGVDMPDDYEDENLYEGLNLDDCSMYEDISRGLQGTYQDVGNLHIGDAQLEKP

Yellow: ITAM

## CD79β-IC

GACAAAGACGATGGTAAAGCCGGAATGGAAGAAGATCATACCTATGAAGGTCTTAATATCGACCA
AACAGCAACCTACGAGGACATCGTAACACTTCGCACAGGCGAAGTGAAATGGAGCGTTGGAGAG
CACCCCGGTCAGGAG

DKDDGKAGMEEDHTYEGLNIDQTATYEDIVTLRTGEVKWSVGEHPGQE
